# An amyloid-β/APP regulatory layer in human brown adipocyte recruitment identifies repurposable drugs for thermogenic fat expansion

**DOI:** 10.64898/2026.10.02.754417

**Authors:** Hannah M. L. Bazin, Sonia Rodríguez-Fdez, Fatima Baldo, Iman D. Mali, Nazuk Gupta, Milidili Maimaiti, Taylor L. Simonian, Mark Campbell, Possawee Prasertsuk, Namshik Han, Antonio Vidal-Puig

## Abstract

Brown adipose tissue (BAT) increases energy expenditure but is scarce in adults, yet most strategies focus on activating existing thermogenic adipocytes rather than recruiting new ones. We integrate human single-nucleus transcriptomics with protein-interaction and knowledge-graph analyses to identify stage-specific, druggable programmes governing brown adipogenesis.

This pipeline resolved four ordered transcriptional states and three transitions, revealing an amyloid-β/APP regulatory layer and nominating APP as a central signalling hub. Most prioritised compounds altered differentiation or metabolic phenotype, several substantially. Phenserine (APP-lowering) and sacubitril (neprilysin inhibitor) promoted differentiation through distinct mechanisms and stages and could be combined sequentially for benefit. Phenserine increased differentiation across three human models and, in mice, expanded BAT mass and increased thermogenic capacity.

Together, these findings reveal targetable control points in human brown adipocyte recruitment, nominate repurposable drugs for BAT expansion, and establish a generalisable framework coupling single-cell transcriptomics with network-based drug repurposing across staged biological systems.

## Introduction

Obesity affects one in eight people worldwide and is a major risk factor for type 2 diabetes, cardiovascular disease, and cancer^1–3^, yet effective and safe therapeutic options remain limited^4–6^. Brown adipose tissue (BAT) has emerged as a promising therapeutic target due to its role in energy expenditure^7^ and systemic metabolic regulation^8^, as well as its potential to counter metabolic disease^9–12^. Unlike white adipocytes, brown adipocytes are rich in mitochondria and uniquely express uncoupling protein 1 (*UCP1*), which enables non-shivering thermogenesis by dissipating energy as heat^7^. In humans, BAT volume inversely correlates with body mass index (BMI)^10^ and associates with improved cardiometabolic health^13,14^. In mice, BAT transplantation enhances energy metabolism and insulin sensitivity, preventing and reversing obesity^11,12^. GLP-1 receptor agonists have transformed pharmacological control of food intake, but they leave energy expenditure, the other side of the energy-balance equation, largely untapped^5,6^; strategies that increase expenditure are therefore a complementary, rather than redundant, addition to current anti-obesity therapy.

Despite this therapeutic promise, pharmacological strategies to activate BAT have largely failed due to poor selectivity, adverse effects, and species differences^15,16^. Additionally, BAT mass declines with age^17^ and its activity is reduced in obesity^18^, limiting the effectiveness of activation-based approaches. Importantly, BAT precursors retain the capacity to differentiate even in metabolic disease^19^. Promoting brown adipocyte differentiation to expand the pool of mature, activation-prone adipocytes offers several advantages: it can enhance the effects of activation at lower input levels and ensures that newly differentiated adipocytes remain under physiological regulation^20^. However, identifying druggable regulators of human brown adipocyte differentiation is challenging due to the complex interplay among cellular signalling pathways and the difficulty in obtaining suitable biological samples. Single-nucleus RNA sequencing (snRNA-seq) has greatly advanced the molecular characterisation of human brown adipose tissue^21,22^ by enabling the study of differentiation processes within the tissue itself rather than relying solely on preadipocyte isolation and *in vitro* differentiation. Nevertheless, transcriptomic approaches alone cannot determine which regulators are most critical within complex signalling networks. Network analysis addresses this limitation by mapping transcriptionally identified proteins onto protein-protein interaction (PPI) networks to pinpoint highly connected regulators that may control differentiation^23^.

Here, we developed and validated a translational pipeline integrating snRNA-seq and two complementary network-based approaches to identify stage-specific regulators of brown adipocyte differentiation and find approved drugs modulating differentiation for therapeutic repurposing. We found that human brown adipocyte differentiation occurs through a graded series of transcriptional states, with metabolic, cytoskeletal, neurogenic and amyloid-processing programmes engaged in a coordinated manner from early stages, the later nominating amyloid precursor protein (APP) as a regulatory hub. Experimental testing confirmed that seven of eight network-prioritised compounds altered brown adipocyte differentiation or metabolic phenotype *in vitro*, five of them substantially. Two of these, phenserine and sacubitril, promoted differentiation via distinct, APP-linked mechanisms and at different differentiation stages, with the lead candidate further validated across multiple human cellular models and *in vivo* mouse systems. Together, these findings establish a validated translational pipeline for anti-obesity drug repurposing, identify a previously unrecognised amyloid-linked regulatory layer in human brown adipocyte recruitment, and demonstrate that stage-specific combination treatment can further enhance differentiation.

## Results

### An integrated snRNA-seq and network-based strategy for stage-specific drug repurposing

We developed an *in silico* to *in vitro* drug repurposing pipeline that integrates snRNA-seq with two complementary network analysis approaches to systematically identify approved drugs that could be repurposed to target brown adipocyte differentiation (Fig. 1). For the PPI-based analysis, we analysed perivascular adipose tissue (PVAT) from the para-aortic area – a small depot surrounding large vessels enriched in brown adipocytes exhibiting canonical thermogenic gene programs^24,25^. PVAT was selected for this analysis because its available snRNA-seq dataset captured intermediate differentiation stages with higher resolution. In parallel, we applied a knowledge graph (KG)-based approach to human supraclavicular BAT snRNA-seq data, the predominant anatomical location of human BAT^26^, and cross-validated our findings against both published and in-house human supraclavicular BAT datasets ^21,22^.

**Figure 1.**
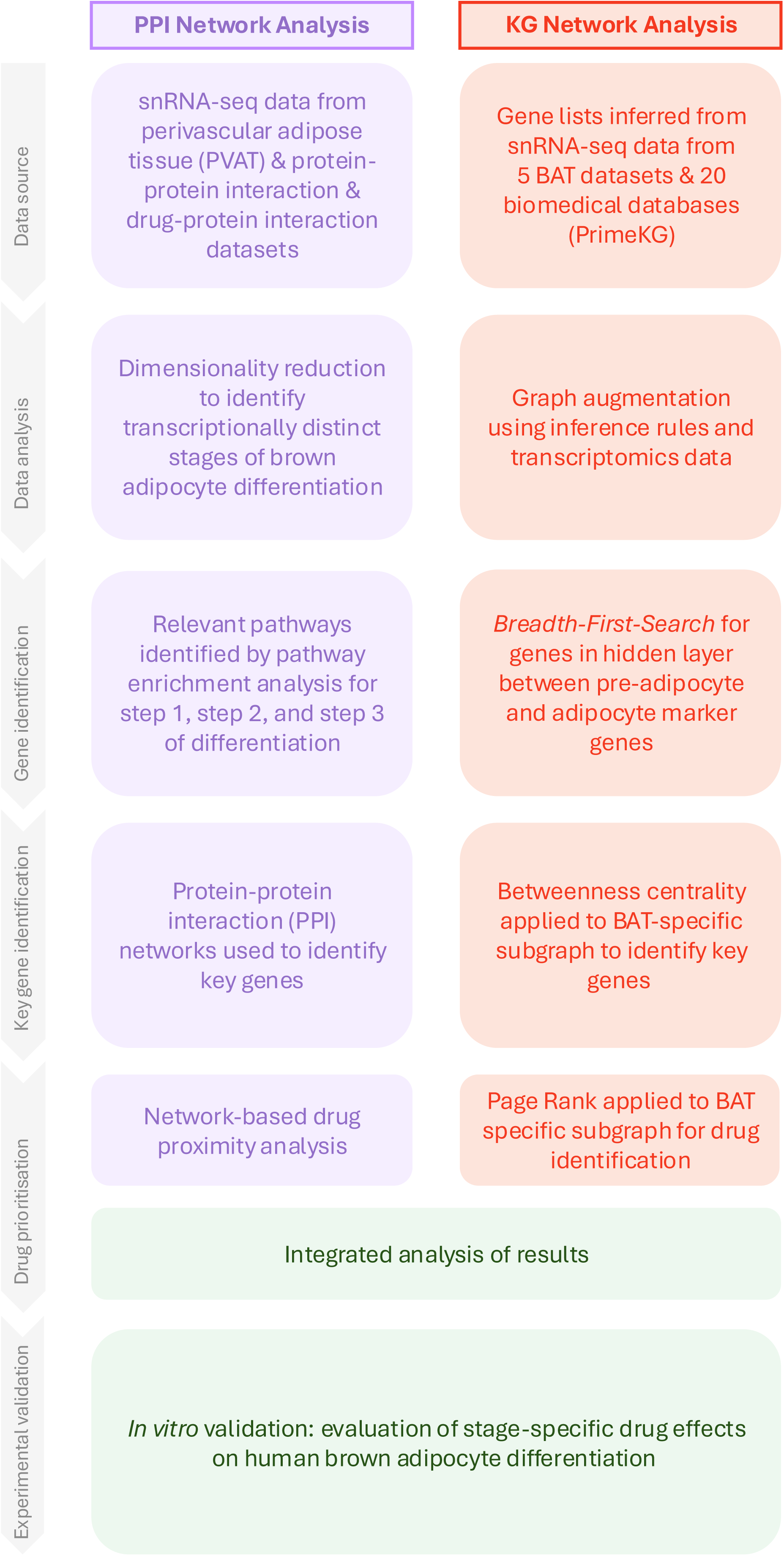
Integrated snRNA-seq and network analysis pipeline for drug repurposing in human brown adipose tissue. Two complementary workflows run in parallel. PPI network analysis (left, purple): snRNA-seq data from perivascular adipose tissue (PVAT), combined with protein-protein and drug-protein interaction data, is used to identify differentiation stages by dimensionality reduction, define stage-specific gene sets by pathway enrichment, identify key genes via PPI networks, and prioritise drugs by network proximity. KG network analysis (right, orange): gene lists from five BAT snRNA-seq datasets and 20 biomedical databases (PrimeKG) are used to augment a knowledge graph, identify hidden-layer genes linking early, intermediate and late differentiation markers by breadth-first search, identify key genes by betweenness centrality, and prioritise drugs by PageRank. Results from both workflows are integrated (green) and candidate drugs validated *in vitro* for stage-specific effects on brown adipocyte differentiation.

Our approach consisted of three integrated steps. First, we identified biological pathways enriched during brown adipocyte differentiation across three discrete stages and the complete trajectory, retaining genes with established roles in adipocyte differentiation or function for the PPI analysis. Second, we applied two independent network-based methods: (1) a PPI network built from these differentiation-associated genes, integrated with a drug– protein interaction database, from which centrality measures identified highly connected regulatory proteins not apparent from expression data alone; and (2), independently, a KG traversal approach identifying regulatory nodes linking preadipocyte- and adipocyte-enriched marker genes, expanded into a BAT-specific subgraph to predict bottleneck genes and their modulating drugs. Third, we prioritised drug candidates from both networks using a network proximity approach^27^, on the premise that drugs whose targets lie closer to key differentiation regulators are more likely to modulate differentiation effectively. The modular design of this pipeline enables application to other cell differentiation systems and disease contexts where single-cell transcriptomic data is available.

### Single-nucleus profiling resolves graded transcriptional stages of brown adipocyte lineage progression

To characterise the transcriptional landscape of brown adipocyte differentiation, we analysed human PVAT snRNA-seq data from three healthy male donors (n=3), providing a cross-sectional view of transcriptional states across the differentiation axis^24^. After quality control filtering, 18,780 high-quality nuclei (75% of the initial 25,197) were retained. Dimensionality reduction and clustering revealed 13 transcriptionally distinct populations (Fig. 2A), including four transcriptional states with progressive brown adipocyte differentiation, hereafter referred to as early preadipocytes, intermediate preadipocytes, transitional adipocytes, and mature adipocytes (Fig. 2B, S1, S2, S3). These states formed a graded differentiation trajectory that we describe as three putative steps. Each state was characterised by distinct marker gene expression (Fig. 2C-D). Early preadipocytes represented the earliest committed adipocyte precursors, characterised by expression of key brown adipogenic priming factors such as *CREB5*, *FSTL1*, *NEGR1*, and *EBF2*^28,29^. Intermediate preadipocytes showed lower expression of *FSTL1* and *CREB5*, alongside higher expression of *MGP*, *ABCA10*, *NEGR1*, and *FBLN1*. Transitional adipocytes were distinguished by the expression of key mature adipocyte markers such as *PPARG*, *ADIPOQ*, *PLIN1*, *PLIN4-5*, *CIDEA*, *CIDEC*, *LPL*, and *GHR*, albeit at lower levels than fully mature cells. Interestingly, this transitional population retains relatively high levels of *EBF2* expression and some immature markers, such as *ITGA11* or *PDGFRA*, although at lower levels than those observed in earlier stages^30,31^. *PPARG* expression, the master transcriptional regulator of adipogenesis, increases throughout differentiation as expected^32^. Mature adipocytes showed high expression of all canonical brown adipocyte markers. We also identified a small *PPARG*+ *EBF2*+ cell population which may reflect the previously described vascular smooth muscle cell-derived brown adipocyte progenitor population^33,34^.

**Figure 2.**
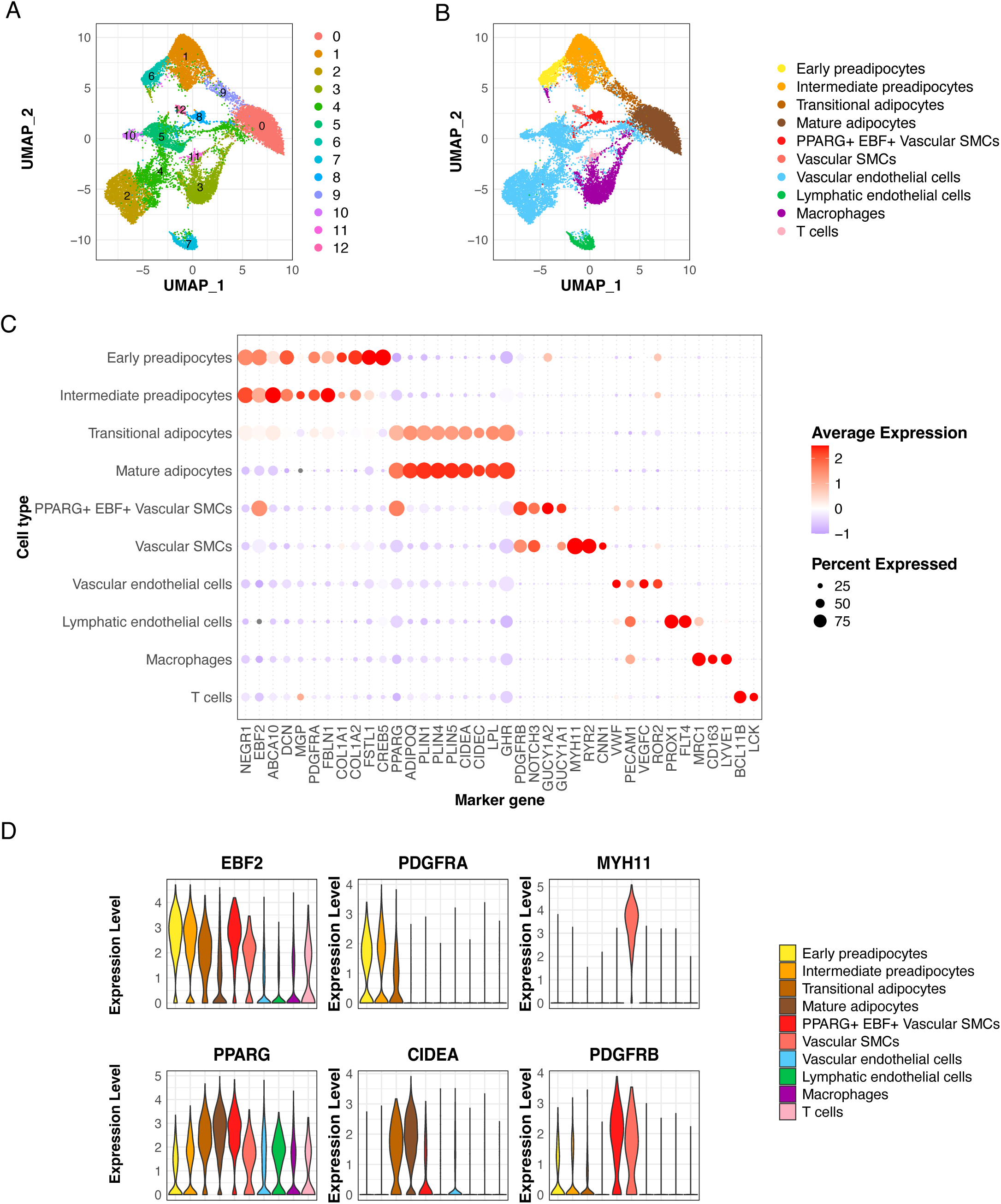
Single-cell transcriptomic analysis identifies four brown adipocyte lineage stages in human perivascular adipose tissue. **(A)** UMAP visualisation of 18,780 nuclei showing 13 distinct clusters. **(B)** Cluster annotation revealing brown adipocyte lineage progression (early preadipocytes, intermediate preadipocytes, transitional adipocytes, mature adipocytes), plus non-adipocyte populations. **(C)** Dot plot showing log-normalised marker gene expression across cell types used to guide cluster annotation. Dot size indicates fraction of cells expressing each gene, colour intensity shows average expression level. **(D)** Violin plots displaying key marker gene expression across cell types. *EBF2* is a known marker of preadipocytes, while *PPARG* expression increases as cells mature. UMAP, Uniform Manifold Approximation and Projection; SMCs, smooth muscle cells.

**Figure 3.**
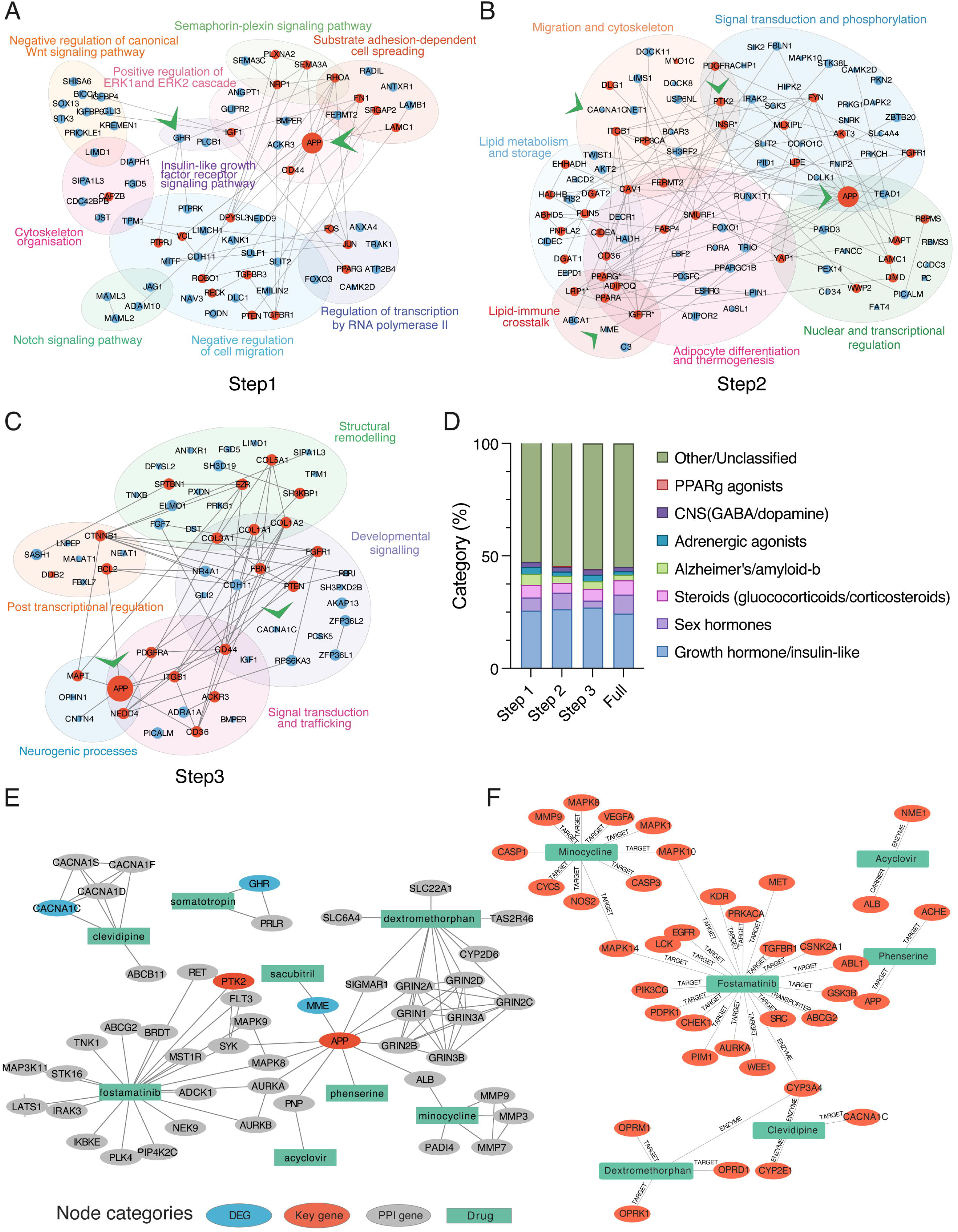
Network analysis identifies eight candidate drugs for *in vitro* validation. **(A-C)** Regulatory modules and candidate nodes for each transition according to PPI analysis **(D)** Main drug categories identified in the PPI-DPI analyses. **(E)** Drug-protein interactions showing eight candidate drugs (green) for modulating brown adipocyte differentiation from PPI-based network proximity analysis. Genes from snRNA-seq analysis are shown in blue, with key genes identified by network centrality analysis in red. PPI genes represent network interactors of DEGs or drug targets; DEGs are differentiation-specific genes. DEG, differentially expressed gene; PPI, protein-protein interaction. **(F)** Drug-protein interaction knowledge graph for KG-based network analysis. The same eight selected drugs are shown with their connected targets.

### Distinct biological processes shape each stage of differentiation

Following cell type identification, pathway enrichment analysis of marker genes revealed biological processes involved in each step of brown adipocyte differentiation (steps 1-3) and in the full differentiation. Beyond canonical adipogenic and metabolic programmes, two recurrent non-canonical themes – neurogenic signalling and amyloid-β processing – emerged across differentiation and were increasingly enriched, prompting us to investigate their potential mechanistic link through network analysis.

Pathway enrichment analysis of the first transition revealed canonical adipogenic processes alongside neurogenic and metabolic priming signatures (Fig. S4). *Roundabout signalling pathway* was the most enriched, with *ROBO1* expression higher in early preadipocytes than in intermediate preadipocytes, whereas *SLIT2* and *SLIT3* were more highly expressed in intermediate preadipocytes. Neurogenic processes related to *axonogenesis* and *axon guidance* were also enriched, involving the ROBO-SLIT signalling axis. These findings suggest that early preadipocytes may already contribute to processes supporting the extensive sympathetic innervation required for BAT activation.

The second transition was dominated by metabolic and lipid-processing pathways, alongside cytoskeletal reorganisation and signalling processes including insulin receptor signalling (Fig. S5)^35,36^. Neurogenic processes remained enriched, consistent with continued interaction with sympathetic innervation. Notably, amyloid-β-related processes emerged at this stage, including amyloid-β clearance, amyloid fibril formation, and cellular responses to amyloid-β.

The final transition was characterised by metabolic and structural maturation, including mitochondrial organisation and actin cytoskeleton remodelling. Neurogenic processes remained prominent and showed progressive enrichment across differentiation. Amyloid-related processes also reached their greatest enrichment at this stage (Fig. S6).

Comparison of early and mature adipocytes confirmed broad metabolic activation, cellular restructuring, and tissue remodelling (Fig. S7), with canonical marker expression following established patterns of brown adipocyte differentiation (Fig. S8)^21,37,38^.

Together, these findings reveal a graded series of transcriptional states, progressing from early cytoskeletal and neurogenic programmes to lipid-processing and, ultimately, mitochondrial and thermogenic maturation. The coordinated enrichment of neurogenic and amyloid-β programmes across differentiation suggests shared regulatory mechanisms, motivating network-based analyses to identify potential hub proteins and druggable targets.

### Orthogonal network-based drug repurposing approaches recover known adipogenic regulators and prioritise novel modulators of brown adipocyte differentiation

To identify therapeutic compounds that might modulate brown adipocyte differentiation, we applied two independent network-based approaches (Fig. 1). The first approach uses a PPI network and a drug-protein interaction (DPI) network, while the second employs a KG. Both methods identified novel drug candidates, with overlapping hits prioritised for experimental validation.

Importantly, these two approaches utilised transcriptomic data from different BAT depots – PVAT for the PPI-based analysis and deep neck BAT for the KG approach – allowing cross-depot validation of our findings. The convergence of results from these anatomically distinct depots strengthens confidence that identified modulators represent broadly applicable therapeutic targets rather than depot-specific regulators.

### PPI-based network analysis recovers known regulators and nominates stage-specific drug candidates

We used a network-based proximity algorithm to integrate PPI and drug-protein interaction (DPI) networks with stage-specific differentially expressed genes (DEGs, Fig. S9)^27^. DEG sets were compiled for each differentiation step and the full differentiation programme and integrated into PPI networks, from which network centrality measures identified key regulatory genes (Fig. 3A–C, Supplementary Tables 5–12).

Drug proximity analysis identified 314–799 significant candidates across the differentiation stages, using either the complete DEG sets or key regulatory genes (Supplementary Tables 13–20). Several therapeutic classes were consistently represented. The analysis recovered established brown adipocyte differentiation activators, including PPARγ agonists and adrenergic receptor agonists (Fig. 3D)^32,39^, supporting the biological relevance of the approach. Hormone-related and central nervous system drugs were also prominent. Notably, drugs associated with amyloid-β clearance and Alzheimer’s disease emerged, including monoclonal antibodies and cholinesterase inhibitors, mirroring the progressive enrichment of amyloid-β processing pathways identified by transcriptomic analysis (Fig. S5-6, Fig. 3A-C).

### KG-based identification further links APP and hormone signalling to brown adipocyte differentiation

To provide an independent perspective on regulatory mechanisms in brown adipocyte differentiation, we applied a KG approach using distinct data sources and analysis methods. First, we constructed a preadipocyte-to-adipocyte network^40^ to investigate previously unrecognised regulatory genes in BAT differentiation. A breadth-first shortest-path search between preadipocyte and mature adipocyte marker nodes identified additional regulatory nodes not apparent from transcriptomic analysis alone, including *MAD2L1*, *PRKCZ*, *CFLAR*, *LIMA1*, *DBN1*, *TMED10*, *ATF3*, and *FN1* (Fig. S10). Several showed stage-dependent expression or phosphorylation patterns consistent with roles in differentiation (Fig. S11), while *AR*, *ESR1*, and *APP* provided additional links to the hormone- and amyloid-related signals identified by the PPI analysis (Fig. 3A-C, Supp. Table 9-16). The resulting network contained 1,975 drug–gene pairs, with hormone receptor modulators, kinase inhibitors, and compounds targeting protein-modification pathways among the prominent drug classes. Together, the KG analysis expanded the regulatory landscape beyond transcriptional changes, identifying pharmacological interactions across multiple regulatory layers, including transcriptional regulation, kinase signalling, protein modification, and protein–protein interactions. This organization supports a view of BAT differentiation as governed by multiple regulatory tiers, with actionable drug targets distributed across gene expression and protein function.

### Integrating PPI and KG results prioritises novel drug candidates for validation

Results from the PPI- and KG-based network analysis (Supp. Table 1, 9-16) were compared to identify convergent candidates. The recovery of well-established regulators, including PPARγ and adrenergic receptor agonists, provided an internal positive control for the biological relevance of the network-based prioritisation. Predefined exclusion criteria removed from this selection compounds targeting established regulators of the differentiation (i.e. PPARγ, PDGFRA regulators), corticosteroids, adrenergic agonists and sex hormones, as well as kinase inhibitors used in cancer treatment whose side effects would likely outweigh their potential metabolic benefits. The remaining candidates were prioritised through an integrated assessment of network proximity scores, established clinical safety profiles, lipophilicity, and mechanistic diversity (Fig. 3E,F; see Methods). The final set was selected to include candidates predicted to affect each differentiation stage of the trajectory and spanning multiple therapeutic classes, including neurotransmitter modulation (dextromethorphan, phenserine), antiviral activity (acyclovir), antibiotic action (minocycline), cardiovascular targets (clevidipine, sacubitril), kinase inhibition (fostamatinib), and hormone signalling (somatropin) (Supp. Table 2). Reassuringly, recruitment conditions in mouse BAT (3 weeks of cold), selectively upregulated the network nodes APP, PTK2 and MME, whereas general adipocyte differentiation markers rose across fat depots (Fig. S12A–H).

### Human *in vitro* screening validates predicted novel stage-specific modulators of brown adipocyte differentiation

To determine whether the eight novel prioritised drugs functionally modulate brown adipocyte recruitment, we tested their effects in human immortalised brown preadipocytes. Drugs were administered at early, intermediate, or throughout differentiation and at multiple concentrations to assess stage- and dose-dependent effects. To maximise the biological information obtained, we implemented a multiparametric high-content confocal imaging pipeline that quantified, on a per-cell basis, the proportion of cells containing lipid droplets, lipid droplet size distributions, and mitochondrial content, allowing combined and individual effects on differentiation, lipid accumulation and mitochondrial content to be distinguished.

Among the prioritised compounds, phenserine and sacubitril emerged as the strongest pro-differentiation candidates, with distinct temporal and phenotypic profiles (Fig. 4, Fig. S13A,B). Phenserine, an Alzheimer’s-associated compound with both acetylcholinesterase inhibitor and APP-downregulating effects^41,42^, increased the proportion of lipid-laden cells only when administered during intermediate stages or throughout differentiation and promoted earlier lipid accumulation (Fig. 4, Fig. S13B,C). At the highest tested dose during intermediate stages, phenserine also reduced lipid droplet size, without affecting mitochondrial content or membrane potential (Fig. 4). Sacubitril, a neprilysin inhibitor, also enhanced brown adipocyte differentiation, but with a different temporal profile. Intermediate doses of sacubitril during the early phase of the differentiation increased both lipid droplet number and mitochondrial mass (Fig. 4, Fig. S13A). Similar results were obtained when cells were treated during the whole differentiation process, but not when treated exclusively during the intermediate stages. At the highest dose, sacubitril also increased lipid droplet size when administered throughout differentiation.

**Figure 4.**
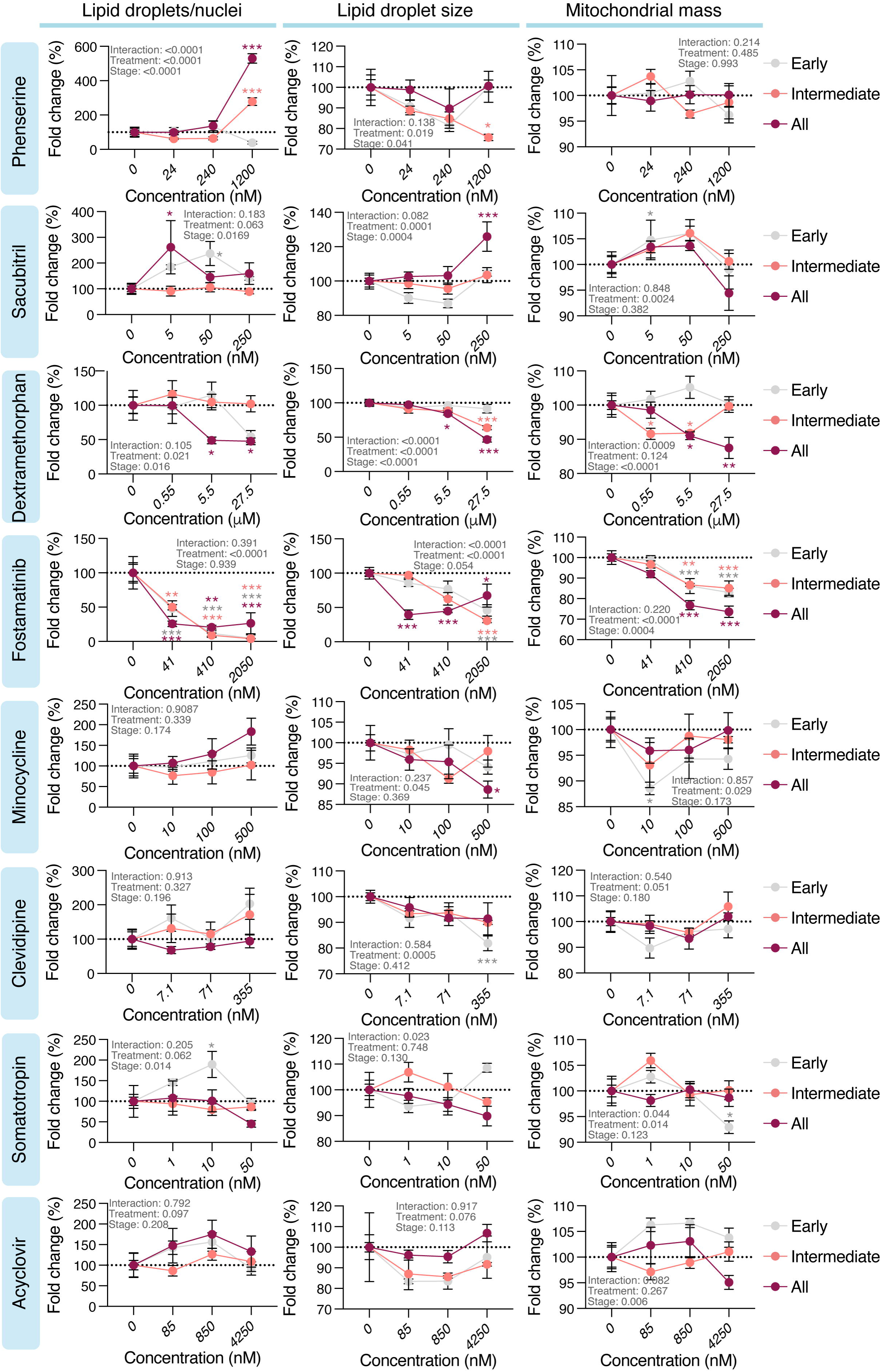
*In vitro* validation reveals the effect of the selected drugs in brown adipocyte differentiation. Human immortalised preadipocytes were differentiated *in vitro* and treated with the indicated compounds and concentrations during the early stages (day 0 to day 6), intermediate (day 6 to day 12) or throughout the whole differentiation process (all). Cells were stained and high-content confocal imaging was used to evaluate different parameters. Two-way ANOVA tested for treatment, stage at which it was added, and interaction effects. Dunnett’s post hoc tests were used for pair comparisons when appropriate. Data represents mean ± SEM with respect to untreated control, with n=4 independently differentiated and treated wells per dose and stage. *, p < 0.05; **, p < 0.01; ***, p < 0.001.

Several compounds showed inhibitory or more selective effects. Fostamatinib and dextromethorphan inhibited differentiation across the measured parameters, although the highest fostamatinib dose was toxic. Minocycline, clevidipine, and somatropin produced more selective effects on lipid or mitochondrial phenotypes, whereas acyclovir had no detectable effect (Fig. 4, Fig. S13C).

### Validated modulators enhance thermogenic and metabolic function in human adipocyte models

To better understand the effects of the drugs with positive effects during the screening, cells were differentiated again in presence of phenserine, sacubitril and minocycline for further functional assessment. Although the effects of minocycline were less pronounced than those of phenserine or sacubitril, it was selected for functional validation due to its connectivity to both compounds in the PPI network (Fig. 4, Fig. S13C, Fig. 3D). Treatment with either minocycline or phenserine increased forskolin-stimulated lipolysis. This was measured by quantification of glycerol release and confirmed at the single-cell level by assessing phosphorylation of hormone-sensitive lipase (HSL) at Ser660 as a readout of the responsiveness and lipolytic capacity of individual cells (Fig. 5A-B). Treatment with minocycline increased the percentage of phosphoHSL positive cells, the average signal intensity, and the lipolysis rate exclusively in response to forskolin but not norepinephrine, suggesting that minocycline acts on signalling pathways downstream of adrenergic receptors rather than at the receptor level (Fig. 5A-B). By contrast, treatment with phenserine had a significant effect on phosphoHSL levels in both untreated and norepinephrine-stimulated conditions (Fig. 5A, Fig. S13B), with higher lipolysis rates when treating with either norepinephrine or forskolin. This beneficial effect also observed for intermediate doses of phenserine in the case of forskolin-treated cells (Fig. 5B). Treatment with sacubitril had a tendency towards increased phosphoHSL levels in basal and forskolin-treated cells, but these small increases were not captured in the less sensitive lipolysis assay (Fig. 5A-B). In line with these results, cells treated with the highest dose of phenserine have increased protein abundance of the PPARG targets HSL, PLIN1 and FABP4 compared to controls, as well as higher levels of phosphorylated HSL (Fig. 5C). Taken together, these results are consistent with both a larger number of differentiated cells and improved individual activation upon phenserine treatment.

**Figure 5.**
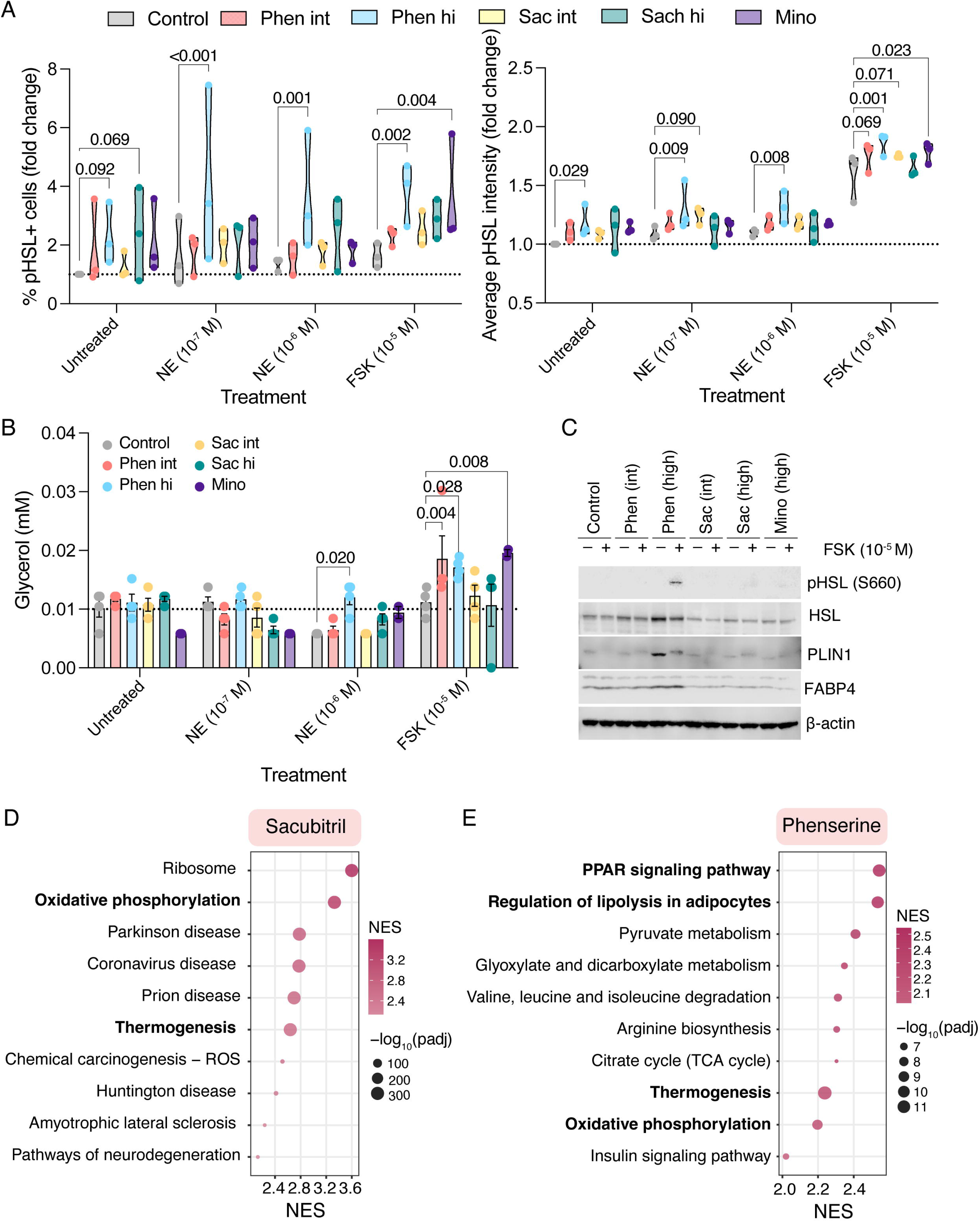
Positive effects translate into improved metabolic responses. **(A)** High-content imaging quantification of phospho-hormone-sensitive lipase (pHSL Ser660) showing mean fluorescence intensity (left) and percentage of positive cells (right). N = 3 independent experiments, each performed in quadruplicate. Repeated-measures two-way ANOVA tested for treatment, stage at which it was added, and interaction effects. Dunnett’s post hoc tests were used for pair comparisons against their relative control. Data are shown as violin plots with the central line representing the median and with respect to untreated control. **(B)** Lipolysis assay in treated versus control conditions. Data shown as mean ± SEM and analysed as above. **(C)** Representative immunoblot analysis of N = 3 independent experiments. For all assays, cells were serum-starved and then treated with the indicated compounds or vehicle control for 90 minutes prior to analysis. **(D,E)** Pathway enrichment analysis on transcriptomic data obtained from fully differentiated cells (day 20) treated with sacubitril in the early stage (D) or phenserine during the intermediate stage (E) of differentiation. NES, normalised enrichment score; padj, adjusted p value.

Although cells treated with minocycline showed significantly higher forskolin-stimulated lipolysis rates, cellular respiration was impaired (Fig. S14A-C). This result is consistent with our initial screening (Fig. 4, right panel) and suggests a complex phenotype where the drug might affect independent signalling pathways during brown adipocyte differentiation. The beneficial effects of the phenserine and sacubitril were further supported by transcriptomic analyses, in which oxidative phosphorylation and thermogenesis were amongst the most enriched pathways (Fig. 5D,E). In the case of phenserine-treated cells, PPAR signalling pathway and lipolysis emerged as the top upregulated pathways (Fig. 5E), confirming our previous observations (Fig. 5A-C). Upstream regulator analysis predicted activation of SREBF1, PPARA, PPARG and SREBF2, all well-known transcriptional regulators of the differentiation process (Fig. S14D).

Sacubitril-treated cells had a different transcriptional profile, identifying APEX1 and HIVEP2 as the most strongly activated transcription factors, together with several known transcriptional regulators, including HSF1, REL and PPARA, suggesting alternative mechanisms underlying the pro-differentiation response (Fig. S14E). Interestingly, principal component analysis further indicated that sacubitril had greater effects at intermediate stages of differentiation, with treated cells progressively converging towards control cells by the end of the process, suggesting that the effects of early treatment are partially attenuated as differentiation progresses (Fig. S14F). This is consistent with the early treatment window identified in the screening (Fig. 4).

In addition to brown adipocyte-related pathways, neuronal and Alzheimer-related pathways were also present amongst the top enriched pathways in the case of sacubitril (Fig. 5D, Fig. S14H), highlighting again the connection shown during the network analysis (Fig. S4-7, Fig. 3A-C).

### Phenserine is a strong promoter of thermogenic differentiation across multiple *in vitro* models and expands BAT *in vivo*

To confirm the translatability of these findings, we assessed the effects of phenserine in two independent human model systems: PAZ6 cells, an SV40 T/t antigen–immortalised human brown preadipocyte line derived from the stromal vascular fraction of infant brown adipose tissue^43,44^ and a hiPSC-based brown adipocyte differentiation platform established and optimised in-house^45^. In both systems, phenserine supplementation enhanced differentiation. In PAZ6 cells, phenserine elicited earlier appearance of lipid droplets in the cells (Fig. 6A), increased basal and FSK-stimulated lipolysis in a dose-dependent manner (Fig. 6B), and increased the phosphorylation of HSL together with UCP1 and FABP4 abundance (Fig. 6C). In the hiPSC model, phenserine was also able to induce earlier expression of *UCP1* (Fig. 6D).

**Figure 6.**
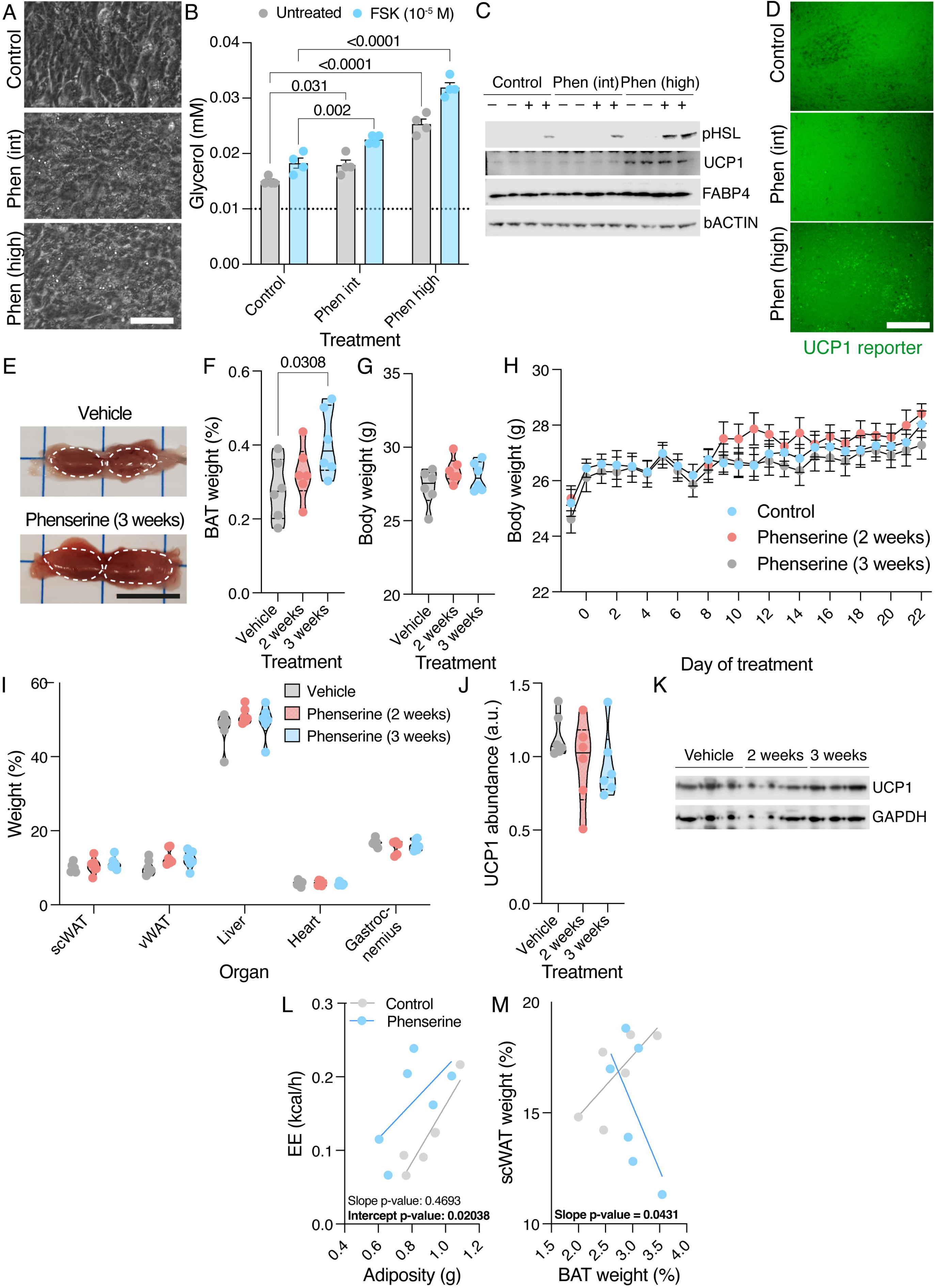
Phenserine promotes brown adipocyte differentiation across different human models and mouse. **(A)** Representative picture of PAZ6 cells on day 9 of the differentiation. **(B)** Lipolysis assay in treated versus control conditions in PAZ6 cells. Cells were starved overnight and stimulated for 90 minutes with or without forskolin before the assay. N=4 wells per condition; phenserine effect confirmed in two additional independent experiments. Data are shown as shown as mean ± SEM. Two-way ANOVA followed by Dunnett’s post hoc tests were used for individual comparisons against their relative control. **(C)** Immunoblot in cells treated with phenserine and stimulated as before. **(D)** Representative picture of UCP1 signal in hiPSC-derived brown adipocytes on day 34 of the differentiation. **(E)** Representative image of the BAT of mice treated with vehicle or phenserine (daily 5 mg/kg intraperitoneal injection) of phenserine for 3 weeks. Scale = 1 cm. **(F)** Weight of BAT at the end of the experiment relative to body weight. **(G)** Total body weight of mice at the end of the experiment and **(H)** body weight evolution. **(I)** Relative weight of the indicated organs at the end of the experiment. **(J-K)** Densitometric quantification (J) and representative immunoblot (K) of UCP1 abundance in the BAT of animals treated with phenserine. **(L-M)** ANCOVA analysis of norepinephrine-stimulated energy expenditure (EE) against adiposity (L) or linear regression of BAT vs scWAT relative weight (M) in vehicle- and phenserine-treated mice after 3 weeks of treatment and 3 days of cold exposure (10°C). In F-J, one-way ANOVA followed by Dunnett’s post hoc tests were used to assess the effect of treatment vs control. In L-M, ANCOVA or linear regression analyses comparing slope and intercept were performed. Data are shown as either violin plots, with the central line indicating the median or as mean ± SEM. N = 6 animals per group.

To assess the therapeutic potential of phenserine *in vivo*, mice were treated daily for two or three weeks with either phenserine or vehicle. Phenserine increased BAT mass (Fig. 6E,F), without differences in final body weight, body weight gain or other analysed organs weight (Fig. 6G-I). UCP1 abundance normalised to β-actin was comparable across groups (Fig. 6J,K), suggesting similar UCP1 levels per unit of tissue but increased total thermogenic capacity as a consequence of BAT expansion. Although no apparent whitening was evident on gross examination (Fig. 6E), resting serum triglycerides but not free fatty acids were elevated (Fig. S15A,B), and BAT vacuole coverage was increased despite conserved nuclei density (Fig. S15C,D); when normalised to BAT weight, estimated total nuclei number showed a non-significant but substantial increase at 3 weeks (Fig. S15E; +30%, p = 0.14), consistent with, though not conclusive for, a contribution from increased adipocyte number alongside hypertrophy. This pattern is atypical for simple hypertrophy alone and consistent with increased acquisition of a lipid-containing phenotype, in line with intermediate-stage *in vitro* effects (Fig. S5, Fig. 4), though a systemic contribution cannot be excluded. Glucose tolerance was unchanged, with phenserine-treated mice showing a non-significant trend toward improved clearance (Fig. S15F). For functional assessment, mice were housed for 3 days at 10°C after 3 weeks of treatment to allow further maturation, and indirect calorimetry performed. Phenserine-treated mice exhibited higher norepinephrine-induced energy expenditure after adjustment for adiposity (Fig. 6L). The expected positive relationship between EE and body weight observed in control mice was not preserved in phenserine-treated mice (Fig. S15G), and the relationship between BAT and WAT mass was likewise altered (Fig. 6M). The scWAT-gWAT relationship remained unchanged (Fig. S15H). Serum triglycerides and free fatty acids were similar under these conditions, in contrast to the resting state (Fig. S15I-J). Together, these findings indicate that phenserine alters BAT composition and enhances maximal thermogenic capacity in lean mice.

### Phenserine acts through a partially APP-dependent, cholinergic-independent mechanism

Phenserine acts through a dual mechanism, both as a direct repressor of APP translation and as an acetylcholinesterase inhibitor^42^ (Fig. 7A). To determine which contributed to enhanced differentiation, we pharmacologically inhibited the muscarinic acetylcholine receptors – those most abundant in brown adipocytes – using two structurally distinct antagonists. None of them was able to inhibit the beneficial effects of phenserine (Fig. 7B,C). Consistent with this, neither the cholinergic agonist carbachol nor the acetylcholinesterase inhibitor rivastigmine (also clinically used in Alzheimer^46^) increased differentiation (Fig. S16A). Carbachol modestly increased phosphoHSL signal intensity at intermediate doses, but less than phenserine, suggesting it might have a modest effect on acute activation (Fig. S16B). This response was not reproduced by rivastigmine, and both treatments decreased mitochondrial mass at high doses (Fig. S16B,C). To further assess specificity relative to other neuroactive agents, we treated cells with memantine, an anti-Alzheimer’s drug targeting NMDA receptors^47^. No significant effects were observed either (Fig. S16A-C).

**Figure 7.**
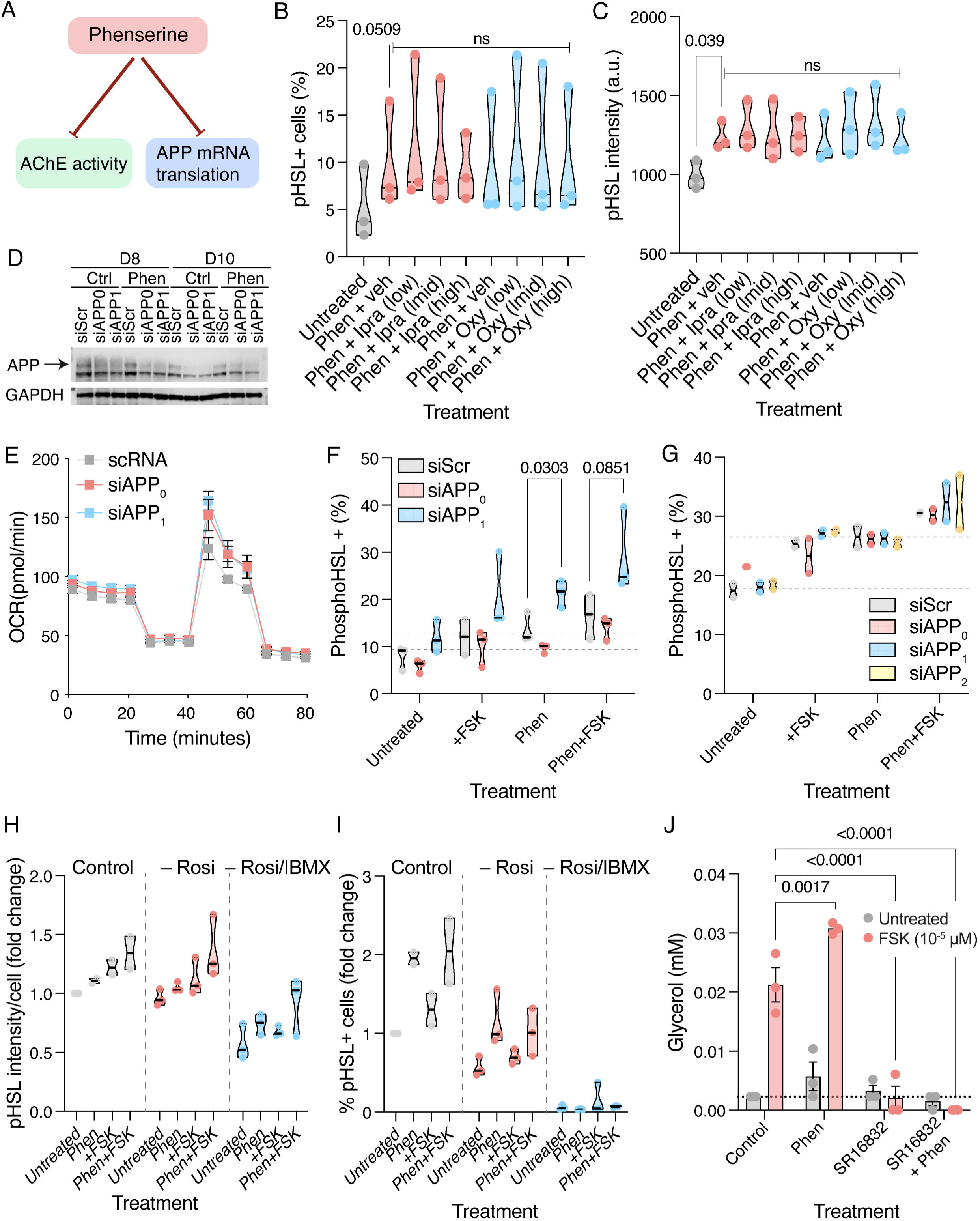
Phenserine promotes differentiation in a cholinergic-independent manner. **(A)** Schematic representation of main axes of phenserine mechanisms of action. **(B-C)** High-content imaging quantification of phospho-hormone-sensitive lipase (pHSL Ser660) showing percentage of positive cells (B) and mean fluorescence intensity (C). N = 3 independent experiments. Repeated-measures two-way ANOVA tested for treatment, stage at which it was added, and interaction effects. Dunnett’s post hoc tests were used for pair comparisons against their relative control. Data are shown as violin plots with the central line representing the median and with respect to untreated control. **(D)** Western blot showing APP silencing. **(E)** Representative Seahorse Mito Stress Test profile from two independent experiments, each performed with two independent siRNAs. **(F-I)** High-content imaging quantification of proportion (F,G,I) and average cell intensity of phospho-HSL (H) in cells where APP was silenced at day 5 of differentiation (F), day 5 and 9 (G) or differentiated in absence or presence of the indicated compounds (H,I). N = 3 independent experiments. Repeated-measures two-way ANOVA tested for treatment, stage at which it was added, and interaction effects. Dunnett’s post hoc tests were used for pair comparisons against their relative control. Data are shown as violin plots with the central line representing the median and with respect to untreated control. Dashed line indicates the values obtained upon phenserine treatment in the same experiments. **(J)** Lipolysis assay in cells differentiated in presence of the PPARg inhibitor SR16832 (2 μM) and or phenserine (12000 nM) from day 6 onwards and stimulated as above. Phen, phenserine; Ipra, ipratropium; oxy, oxybutinin; Rosi, rosiglitazone; scRNA, scramble siRNA; IBMX, 3-isobutyl-1-methylxanthine.

To directly test the contribution of APP to the phenotype observed, we silenced the gene on day 5, coinciding with the window during which phenserine treatment was most effective (Fig. 4, Fig. 7D). *APP* downregulation using two independent siRNAs increased maximal respiration, supporting a role for APP-linked signalling in brown adipocyte metabolic maturation (Fig. 7E). However, APP silencing did not fully recapitulate the effects of phenserine, even when performed twice during the responsive window. Moreover, treatment with phenserine elicited comparable increases in the percentage of phosphoHSL positive cells in control and APP-silenced cells, suggesting that APP contributes but not fully mediates the phenserine response (Fig. 7F,G). Together with the lack of evidence for a cholinergic mechanism, these data position APP as a network-nominated entry point into a broader phenserine-responsive differentiation programme. Consistent with this interpretation, transcriptomic analysis of phenserine-treated cells showed a strong activation of the master transcriptional regulators PPARG and SREBF1 (Fig. 5C,E, Fig. S16D), suggesting convergence on brown adipocyte maturation programmes.

Finally, phenserine retained its pro-differentiation effects in the absence of the exogenous PPARγ agonist rosiglitazone and under reduced adipogenic induction conditions (no rosiglitazone or IBMX), although fewer cells reached a fully differentiated state (Fig. 7H,I). Conversely, pharmacological inhibition of PPARγ blocked differentiation and lipolysis irrespective of phenserine treatment (Fig. 7J). These results indicate that phenserine acts within an intact PPARγ-dependent adipogenic framework, rather than acting simply as a substitute for exogenous PPARγ agonism.

### Data-informed stage-specific drug combinations enhance differentiation

After identifying and characterising stage-specific modulators, we next wanted to determine whether the combination of compounds acting at distinct stages of the differentiation could be combined to further enhance brown adipocyte differentiation. Motivated by the observation that the maximal effects of early-stage modulators could be at intermediate phases (Fig. S14F), cells were treated with the early-stage modulator sacubitril during the initial part of the differentiation, either as a single agent or followed by co-treatment with the intermediate-stage modulator phenserine during the intermediate phase (Fig. 8A). This sequential treatment increased uncoupled respiration, as reflected by elevated proton leak (Fig. 8B,C). High-content imaging analysis revealed comparable increases in both average phoshoHSL intensity and proportion of phosphoHSL positive cells using the combination regimen and phenserine alone (Fig. S14A,B). However, immunoblot analysis revealed a markedly greater induction of phosphoHSL upon stimulation, together with increased abundance of HSL, PLIN1, FABP4 and some mitochondrial complexes consistent with an enhanced population-level response (Fig. 8D).

**Figure 8.**
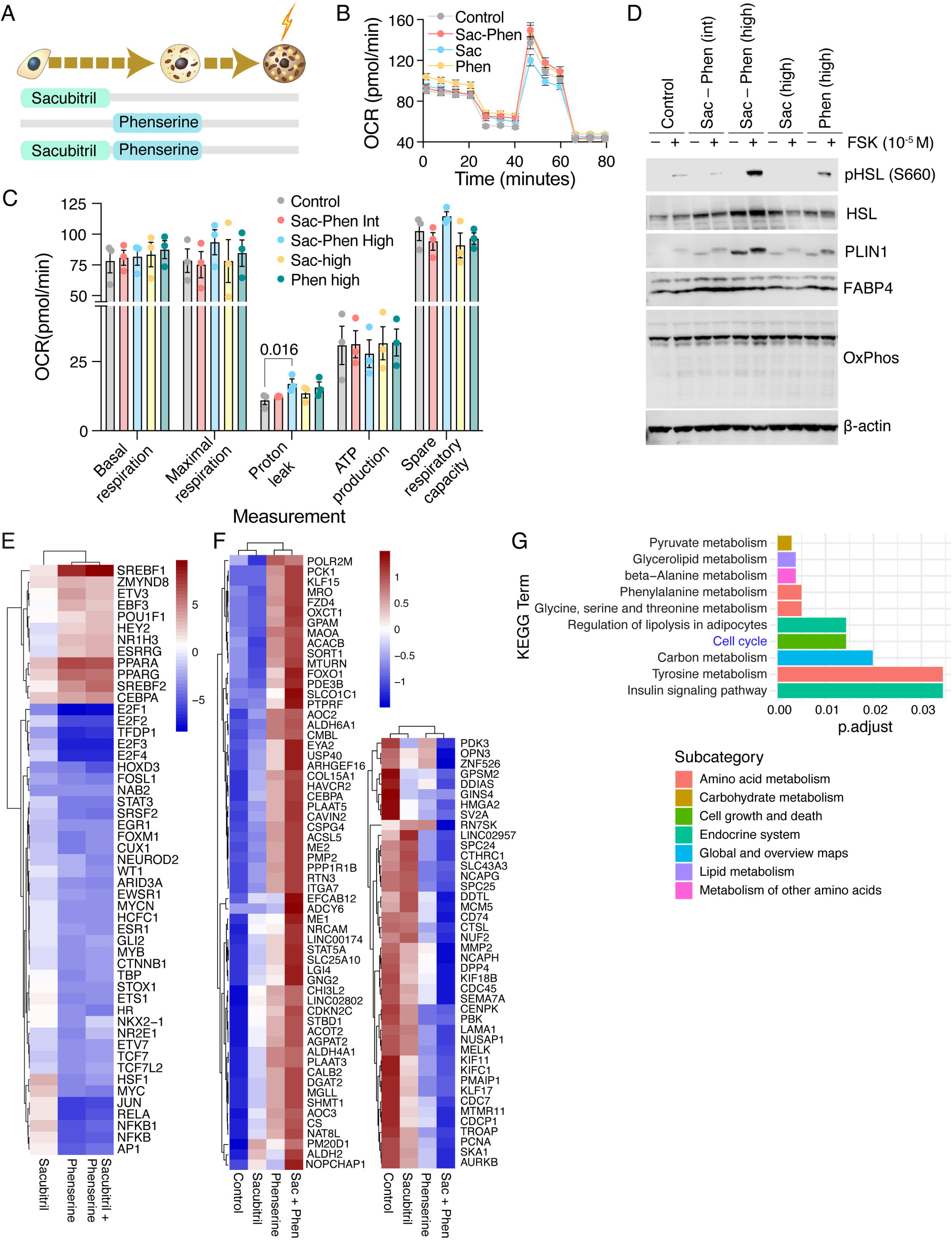
Combination therapy shows beneficial effects over the use of individual drugs. **(A)** Schematic representation of the treatments. **(B-C)** Representative Seahorse Mito Stress Test profile (B) and quantification in differentiated hTERT-BA cells treated with the indicated compounds and concentrations throughout the differentiation process. Repeated-measures two-way ANOVA was used to assess treatment, dose, and treatment × dose interaction effects, followed by Dunnett’s post hoc test comparing each condition with the corresponding untreated control. Data are presented as mean ± SEM from three independent experiments and are normalized to the untreated control. **(D)** Representative immunoblot of mature cells differentiated in presence of the indicated drugs and combinations. **(E)** Heatmap representation of significantly altered transcription factor activity as inferred from transcriptomic data upon the indicated treatments. **(F,G)** Heatmap of normalised gene expression (F) and pathway overrepresentation analysis (G) of top supra-additive genes in the dual treatment vs the expected combined effect of each single drug. N = 3 independent experiments per condition.

Transcriptomic analysis revealed broadly similar patterns of transcription factor activation in cells treated with phenserine alone or in combination, although the sequential treatment elicited a more robust activation of SREBF1, SREBF2 and CEBPA amongst others (Fig. 8E). Analysis of supra-additive or combination-enhanced genes, defined as genes whose expression changes exceeded the sum of those induced by the individual treatments, identified enhanced upregulation of pathways related to insulin signalling, lipolysis, and lipid and protein metabolism, accompanied by downregulation of cell-cycle programmes (Fig. 8F,G), demonstrating that compounds acting at distinct stages of differentiation can be rationally combined to enhance brown adipocyte maturation.

## Discussion

Incretin-based therapies have transformed obesity treatment by acting on energy intake. Yet, tolerability, long-term adherence, concomitant lean-mass loss, and rapid weight regain after discontinuation highlight the need for complementary approaches^52,53^. Thermogenic activation increases energy expenditure independently of appetite suppression and could, in principle, help preserve lean mass during weight loss and sustain energy expenditure after GLP-1 discontinuation. Pharmacologically enhancing endogenous recruitment to expand the pool of mature thermogenic adipocytes could help overcome reduced BAT abundance in obese and diabetic individuals and sustain metabolic benefits during BAT activation.

Our findings reveal that human brown adipocyte differentiation involves a series of coordinated transcriptional stages characterised by cytoskeletal reorganisation, metabolic priming and acquisition of the molecular machinery required to store triglycerides in lipid droplets and provide thermogenic substrates^48^ ahead of the mitochondrial remodelling required to enable the non-shivering thermogenic capacity^49^. The specific sequence of this transcriptional programme enables the identification of more directed stage-specific interventions to promote differentiation. We also observed progressive enrichment of neurogenic and synaptic pathways throughout differentiation, consistent with the close relationship between brown adipocytes and sympathetic innervation^50–52^. In particular, the early enrichment of Roundabout signalling^53–55^ during the differentiation point to a local signalling axis and suggest that early differentiating adipocytes may already contribute to the establishment of the neurovascular environment required to support BAT activation.

We selected eight candidate compounds for *in vitro* testing, spanning endocrine (somatropin), natriuretic peptide (sacubitril), kinase (fostamatinib), calcium-channel (clevidipine) and neuropharmacological (dextromethorphan, phenserine) mechanisms. Seven altered at least one differentiation or metabolic readout, and five did so substantially. The two inhibitory hits, fostamatinib and dextromethorphan, are consistent with established roles for their primary targets, SYK and SigmaR1, in BAT development and mitochondrial function respectively ^56,57^, a direction that, while counterintuitive for an anti-obesity strategy, could be relevant to context-dependent suppression of recruitment in sustained hyperadrenergic states such as pheochromocytoma or major burns. Somatropin, previously reported to induce browning ^58^, produced only selective effects on lipid and mitochondrial phenotypes in our system, without enhancing differentiation.

Two drugs promoted differentiation in a stage-specific manner. Sacubitril, a neprilysin inhibitor, emerged as an early-stage modulator. Its effects may reflect increased natriuretic peptide availability, including atrial natriuretic peptide (ANP), previously linked to browning^59–61^. Neprilysin also contributes to the degradation of APP-derived peptides, including amyloid-β, providing a potential link between sacubitril treatment and the APP- and Alzheimer’s disease-related pathways identified in our transcriptomic analysis^62^. Given the heart’s role as a major source of ANP, this mechanism may also be relevant to PVAT, where brown adipocyte function may have a cardioprotective role^20,63^.

Phenserine, which has been reported to modulate both acetylcholinesterase activity and amyloid-β processing^64^, enhanced brown adipocyte differentiation across three independent human *in vitro* models and expanded BAT mass in mice. Several aspects of our data argue against a predominantly cholinergic mechanism in this setting: carbachol and rivastigmine did not reproduce the phenotype, and muscarinic antagonism did not attenuate the effects of phenserine. APP knockdown phenocopied some, but not all, phenserine-associated responses, positioning APP as a contributing regulatory node rather than the sole mechanism. Instead, phenserine appeared to engage a broader pro-adipogenic programme, supported by activation of PPAR- and SREBF-related transcriptional regulators and by the requirement for an intact PPARγ-dependent differentiation framework. Thus, phenserine may act through APP-associated and APP-independent mechanisms that converge on brown adipocyte maturation. *In vivo*, beyond increasing BAT mass, the elevation in maximal thermogenic capacity and compositional findings suggest genuine recruitment of thermogenic capacity rather than passive tissue accumulation. The increased vacuole percentage with conserved UCP1 and nuclei density in resting conditions may reflect an intermediate differentiation state in which lipid-handling machinery outpaces mitochondrial maturation, consistent with the ordered human trajectory and *in vitro* data. There could also be a homeostatic regulation compensating for the increased BAT mass masking the added capacity at rest. We cannot formally exclude an organismal contribution to the resting phenotype, but reproduction of phenserine’s effects across three independent human *in vitro* models argues for a genuine cell-intrinsic component.

More broadly, the enrichment of amyloid-related processing and clearance signatures during differentiation, phenserine’s activity in our system and prior observations linking APP-related pathways to BAT biology, including our own finding that the soluble form of the regulator of APP endocytic trafficking SORL1 can modulate thermogenesis^65^, support a shared regulatory axis between BAT and neurodegenerative disease. Consistent with a shared axis, a classic transgenic Alzheimer’s mouse model (Tg2576) carrying a deleterious APP mutation has been shown to exhibit markedly reduced BAT glucose uptake, although BAT itself was not characterised in that study^66^. Phenserine’s own clinical development in Alzheimer’s disease is discontinued^67^ in favor of its next-generation (+)-enantiomer buntanetap (posiphen), which lacks acetylcholinesterase inhibitory activity and its associated side effects and is now in late-stage clinical trials.

The pipeline’s predictive performance is itself a result. Target-led screening would have been unlikely to place an Alzheimer’s compound and a heart-failure drug on a list of candidate brown adipocyte modulators, and both were nominated before any cell was treated. By integrating insights from single-cell transcriptomics with dual network-based drug repurposing approaches, the pipeline reveals how drugs can influence distant biological processes through interconnected pathways, capturing the complexity of cellular systems that single-target approaches cannot address. Dissecting differentiation into distinct stages further allows the identification of stage-specific drugs that can fine-tune this process, potentially enabling individualised approaches based on the predominant precursor subsets present; the distinct, stage-restricted effects of phenserine and sacubitril, and their potentiation when combined sequentially, underscore that timing can be as important as the target engaged. In summary, this validated translational pipeline nominates therapeutic candidates for BAT expansion with potential anti-obesity implications and establishes a broadly applicable strategy for drug discovery across diverse staged or ordered biological systems.

## Methods

Data was analysed in `R` (v.4.4.1)^68^ using RStudio^69^, and in `Python` (v3.9.12) with Visual Studio Code (v1.101.0). Figures were plotted with the `R` package `ggplot2` (v3.5.1)^70^ and assembled with the `R` package `patchwork` (v1.3.0)^71^.

### Single-cell transcriptomic analysis

Data was processed using the `R` package `Seurat` (v4.4.0)^72^. Data was processed following the standard `Seurat` pipeline. Nuclei were filtered using: 1000 < `nFeature_RNA` < 6000, 500 < `nCount_RNA` < 40000, and `percent.mt` < 1, resulting in three samples with 9,420, 4,390 and 4,970 cells. Gene expression was normalised and variable features identified. To minimise batch effects, the three objects were integrated and scaled, regressing out mitochondrial content. Fifteen dimensions were selected for downstream analysis. Thirteen clusters were identified with `FindNeighbors` and `FindClusters` (`resolution=0.35`). Differential gene expression analysis was performed on the RNA assay using `FindAllMarkers` to identify cluster-specific markers (`min.pct=0.3`, `logfc.threshold=0.3`). Cluster annotation was guided by marker gene expression and through visualising predefined canonical markers.

### Pathway enrichment analysis

Pathway enrichment analysis was performed using the `R` package `pathfindR` (v2.4.1)^73^. The integrated `Seurat` object was filtered to retain adipocyte lineage clusters only (early preadipocytes, intermediate preadipocytes, transitional adipocytes, mature adipocytes). Scaling was recomputed, regressing out mitochondrial content. DEGs were identified for differentiation steps 1-3 (step 1: early vs intermediate, step 2: intermediate vs transitional, step 3: transitional vs mature) and full differentiation (early vs mature) using `FindMarkers` (`min.pct=0.3`, `logfc.threshold=0.3`) from `Seurat`. For each DEG list, pathway enrichment was performed using `run_pathfindR` (`gene_sets="GO-BP"`, `min_gset_size=5`, `max_gset_size=300`) from `pathfindR`. After filtering (lowest_*p*<0.05, `occurrence>5`, i.e. enrichment frequency over 10 iterations), the top 20 pathways by fold enrichment were manually grouped into themes and visualised. Heatmaps were generated using `DoHeatmap` from `Seurat` on the RNA assay for marker gene expression across cells, `AverageExpression` from `Seurat` for average gene expression across clusters, and `pheatmap` (v.1.0.12)^74^ for pathway fold enrichment.

### PPI-based network analysis

To identify and prioritise drug targets and compounds, a network-based algorithm by Han et al.^27^ was used, requiring as inputs a PPI network, a DPI network, and a gene list. This analysis was performed in `Python` using `NetworkX` (v5)^75^. A full list of dependencies is available in the `requirements.txt` file in the project repository (see Data Availability). Certain steps were performed in `R` using `biomaRt` (v2.60.1, Ensembl v114 GRCh38.p14, accessed May 2025)^76,77^.

### PPI network assembly

The PPI list was assembled by combining data from STRING (v12.0, July 2023)^78^, BioGRID (v4.4.245, 30 April 2025) ^79^, and HINT (June 2024)^80^. Ensembl protein IDs were converted to HGNC symbols using `biomaRt`. Filtering was performed to retain medium confidence (≥0.4) human protein pairs (STRING) and multi-validated physical human interactions (BioGRID). All three datasets were merged and duplicates removed. The final list was converted into a network (345,547 edges, 18,451 nodes) (Supplementary Table 3) using `NetworkX`^75^.

### DPI list assembly

The DPI list (drug–target gene interactions) was assembled from DrugBank (19 May 2025)^81^ and ChEMBL (v35, December 2024)^82^. Filters were applied to retain active, human, and high-confidence interactions involving approved or clinically tested drugs. Gene symbols were mapped using a ChEMBL-UniProt mapping file (v35) and `biomaRt`. The final list contained 15,951 unique DPIs (2,249 unique drugs, 1,933 unique targets). Filtering to keep only DPIs with a target in the PPI network reduced it to 15,744 unique DPIs (2,244 unique drugs, 1,903 unique targets) (Supplementary Table 4).

### Differentiation gene list assembly and key gene identification

We reviewed pathway enrichment analysis results and selected for network analysis pathways containing ≥1 known adipocyte regulator and potential regulatory genes. For each step, constituent genes (up- and downregulated) of all selected pathways formed the gene list corresponding to that step. DEG compilation yielded 70, 99, 56, and 168 genes for differentiation steps 1-3 and full differentiation, respectively (Supplementary Tables 5-8). To identify key regulators in each list, network centrality measures were used, following Han et al.^27^. Degree centrality was used to find highly connected nodes, betweenness centrality to identify potential bottlenecks, eigenvector centrality to determine influential proteins, and personalised PageRank to assess how closely each gene’s product is connected to other DEG products. To generate a null distribution, 100 random gene sets of equal size were sampled from the PPI network and their centrality scores calculated. Key genes were defined as having at least one *p*<0.05 across centrality measure. Network centrality analysis (*p*<0.05) identified 22, 37, 21, and 52 key genes per stage (Supplementary Tables 9-12).

### Drug proximity analysis

The PPI network, DPI list and gene lists were used as inputs for the Han et al.^27^ network-based proximity algorithm, which computes the average shortest path length from drug targets to input genes within the PPI network. To determine statistical significance, 1,000 randomised iterations were computed, each time with nodes with matched degree (i.e. number of connections) randomly selected from degree bins. This enabled each drug to receive a *z*-score. A more negative *z*-score indicated closer proximity between the drug and the gene list than expected by chance, suggesting biological relevance. Further details can be found in Han et al.^27^. For each gene list (steps 1-3 and full differentiation), the algorithm was run twice using (1) the full gene list and (2) the key genes only. Drugs with a *z*-score ≤ -2.576 were considered significant (0.01 significance level; calculated with `scipy.stats` v1.13.1^83^). Drug proximity analysis of complete DEG sets identified 641, 779, 636, and 799 significant compounds (*z*≤-2.576, *p*≤0.01) targeting 974, 1213, 1086, and 1050 proteins, respectively (Supplementary Tables 13-16). Analysis restricted to key genes yielded 335, 450, 314, and 518 significant drugs targeting 227, 499, 179, and 364 proteins, respectively (Supplementary Tables 17-20). Mechanism of action information was retrieved from DrugBank and ChEMBL. To prioritise significant drugs for *in vitro* validation, several filters were applied. Drugs with network distance ≤1 were retained, excluding sex-hormone related compounds, adrenergic agonists, glucocorticoids, cancer drugs, and PPARγ/PDGFRA modulators. Prioritisation was further guided by: (1) target gene relevance, (2) confirmation in independent network-based prioritisation, (3) drug lipophilicity, and (4) experimental availability. Visualisation of PPI and DPI networks was done with Cytoscape (v3.10.3)^84^. Drugs chosen for experimental validation and their respective targets are in Supplementary Table 21.

### Knowledge graph analysis

*Data*. PrimeKG^85^ is a public knowledge graph containing biomedical information curated from 20 datasets and harmonised to provide a heterogeneous, undirected biomedical network. It comprises 129,375 nodes and 4,050,249 edges. To enhance connectivity between genes and diseases, we augmented the graph using reference-based logical rules. Specifically, if Protein 1 interacts with Protein 2 via a PPI, Protein 2 is associated with Disease X, and both proteins are expressed in the same anatomical location, we added an edge between Protein 1 and Disease X labelled "ANATOMY_ONE_HOP_RELATED". This augmentation resulted in the addition of 4,900,476 edges to the graph.

### Identification of novel gene regulators of differentiation

To identify genes that may regulate the differentiation process between preadipocyte and adipocyte stages, we imported the graph into Neo4j (Neo4j Technology Inc., 2015), a database management system that stores data as nodes, edges and their associated attributes. We then applied a depth-first search algorithm to trace connections from an initial set of genes–representing pre-adipocyte markers of differentiation (DEGs)–to a second set of genes corresponding to adipocyte differentiation markers. Intermediate nodes encountered along these paths were extracted as a "hidden layer" of genes potentially involved in regulatory processes governing cellular differentiation.

### Drug proximity scoring to BAT-relevant key nodes in the BAT subgraph

To prioritise candidate drug modulators of the regulatory genes, we constructed a BAT-specific differentiation subgraph. Human BAT snRNA-seq data^40^ was reanalysed and differential expression analysis was performed comparing preadipocytes or adipocytes against each of the non-adipocyte cell types. Genes showing >1.5 fold upregulation across all comparisons were considered as brown preadipocyte or adipocyte-enriched, respectively. DEGs for preadipocytes and mature adipocytes were integrated into the graph and breadth-first shortest path algorithm applied to identify connections from DEGs to drug nodes. The algorithm traversed up to two hops from the DEGs, capturing all intervening node types and edges whilst excluding "contraindication", "side_effect", "expression_absent" and "phenotype_absent" edges. The resulting subgraph contained 7,690 nodes and 172,155 edges (Supplementary Table 22). Centrality-based ranking algorithms, including betweenness centrality and PageRank, were then applied to identify key genes likely to influence differentiation processes and their modulators. To identify drugs targeting critical regulatory bottlenecks, suggesting strategic intervention points, we first identified drugs with high PageRank (using a damping factor of 0.85 to ensure convergence and a maximum of 1,000 iterations; ≥90th percentile) connected to genes with high betweenness centrality (≥90th percentile). This query returned 1,975 drug–gene pairs, focusing on drugs as hubs and genes as bottlenecks.

### Cell culture

Human immortalised brown preadipocytes (ATCC, CRL-4062) were cultured and differentiated following supplier’s protocol in a 37°C and 5% CO2 humidified incubator. PAZ6 were cultured as previously described^86^. The UCP1-reporter cell line of pluripotent stem cells (Kolf-C2) was generated by CellSystems (Germany) and cultured as in Stem Cell Reports, 2021^45^. In brief, cells were maintained in TeSR-E8 media (Stem Cell Technologies) and seeded on Matrigel-coated plates for differentiation. Media composition is indicated in Supplementary Table 23. In all cases, media was changed every other day during the differentiation.

### Cell treatment and stimulation

Cells were treated with the indicated drugs added in combination with the regular cytokines and media changed every other day. For stimulation, cells were starved overnight and treated with either norepinephrine or forskolin for 90 minutes prior to fixation or functional assessment. Drugs used are indicated in Supplementary Table 24.

### Cell imaging and staining

PhenoPlate™ 96-well microplates (Cat #6055300, Revvity) were used for imaging experiments. After 16-20 days, cells were gently washed with DPBS, stained with Mitotracker (1:2000, Cat. #M22425, Thermo Fisher Scientific) and LipidTOX Deep Red (1:500, Cat. #H34477, Thermo Fisher Scientific) for 20 min, washed, fixed (4% PFA, 10 min) and incubated with DAPI (1µg/mL) for 5 min before imaging using *Opera Phenix Plus HCS*. For phospho-HSL staining, cells were incubated overnight at 4°C with primary antibody (Cat# 4126, Cell Signaling Technology) diluted in 10% normal donkey serum (Cat. #017-000-121, Jackson ImmunoResearch) in TBS-T. Following three washes, cells were incubated with an Alexa Fluor-conjugated secondary antibody (1:500, diluted as before) for 1 hour, washed, and stained with LipidTOX Deep Red (Cat. #H34477, Thermo Fisher Scientific; 1:1000) and DAPI (1µg/mL) diluted in TBS-T for 30 min in the dark prior to imaging.

### Image acquisition and analysis

Cell plates were imaged in an Opera Phenix High-Content Screening System (Revvity), taking pictures from 13 different fields per well. Images were analysed using Harmony v4.8 software (Revvity). In brief, nuclei were segmented based on the DAPI signal and used to normalise results by number of cells. Fields with less than 50 nuclei were not included in the final analysis. Lipid droplets were analysed as “spots” and a roundness threshold of ≥0.7 was established to calculate the number, size and distribution of lipid droplets. Where phospohHSL staining was present, cell segmentation was performed based on this metric instead. Two-way ANOVA was used to test the effect of concentration and time of treatment of each of the measured parameters, followed by Dunnett’s post hoc test when appropriate.

### Lipolysis assays

Cells were starved overnight in phenol red-free DMEM/F12 medium (Cat. #11039021, Thermo Fisher Scientific) and then either left untreated or treated for 90 min as indicated above. Cell culture supernatants were collected, and glycerol release was measured as proxy of lipolysis using the Free Glycerol Reagent (Cat. #F6428, Sigma-Aldrich), following the manufacturer’s instructions. A glycerol standard curve was generated in parallel for quantification.

### Protein extraction and Western blot

Cell protein was extracted using RIPA buffer (10 mM Tris-HCL pH 7.4, 150mM NaCl, 1% Triton-X-100 and Pierce phosphatase and protease inhibitors (Cat #A32961, Thermo Fisher Scientific). Supernatants were collected, and protein concentration was determined using the Precision Red Advanced Protein Assay (Cat. #ADV02A, Cytoskeleton) according to the manufacturer’s instructions. Equal amounts of protein lysate were loaded onto precast polyacrylamide gels (Cat. #WG1401BOX, Thermo Fisher Scientific) and electrophoresed in MOPS running buffer (Cat. #NP0001, Thermo Fisher Scientific) at constant voltage. Proteins were transferred onto nitrocellulose membranes using the iBlot 2 system (Cat. #IB23001, Thermo Fisher Scientific).

Membranes were blocked in 5% BSA in TBS-T for at least 30 min and incubated overnight at 4°C with primary antibodies (Supplementary Table 25). Following three washes in TBS-T, membranes were incubated for 1 h at room temperature with HRP-linked secondary antibodies (Cell Signaling Technology, Cat. #7074 or #7076; 1:3000 dilution in TBS-T containing 5% BSA). After three further washes in TBS-T, membranes were developed using Immobilon Forte Western HRP Substrate (Cat. #WBLUF0500, MilliporeSigma) and imaged using a LI-COR Odyssey Fc Imaging System (Cat. #2800, LI-COR Biosciences).

### Seahorse analysis

Cells were detached on day 14 using TrypLE Express Enzyme (Cat. #12604013, Thermo Fisher Scientific) and seeded onto Seahorse XF96 cell culture microplates (Cat. #101085-004, Agilent). On days 18–20, cells were serum-starved and treated as appropriate prior to analysis using an XFe96 Seahorse Extracellular Flux Analyzer (Cat. #XFe96, Agilent) with the Seahorse XF Cell Mito Stress Test protocol. Compounds were injected sequentially as follows: oligomycin (5 μM; Cat. #103015-100, Agilent), carbonyl cyanide 4-(trifluoromethoxy) phenylhydrazone (FCCP; 6 μM; Cat. #103015-100, Agilent), rotenone (2 μM; Cat. #103015-100, Agilent), and antimycin A (2 μM; Cat. #103015-100, Agilent). Nonresponding wells were excluded and calculations were performed according to the suppliers’ recommendations. Following the assay, plates were stained with Hoechst 33342 (1:10,000, 15 min; Cat. #62249, Thermo Fisher Scientific), and fluorescence was measured using a Tecan Infinite M1000 Pro plate reader (Cat. #30050303, Tecan) to confirm consistent cell number across wells and treatment groups.

### Animals and treatments

Male C57BL/6J mice were obtained from Charles River. Animals were weighed on arrival and randomly allocated to treatment groups such that body-weight distributions were comparable between groups. 12-week-old mice were treated with either vehicle (1% DMSO in 0.9% saline) or phenserine (5 mg/kg, intraperitoneally, once daily) for 2 (14 days) or 3 weeks (23-24 days). All procedures were carried out under the Animals (Scientific Procedures) Act 1986 Amendment Regulations 2012, following ethical review by the University of Cambridge Animal Welfare and Ethical Review Body (AWERB), in accordance with UK Home Office guidelines. Mice were maintained under specific pathogen-free conditions at 22 °C, 55% relative humidity, and a 12-h light/dark cycle, with ad libitum access to food and water.

### Histological analysis

BAT samples were fixed in 4% paraformaldehyde for 24 h, paraffin-embedded, sectioned at 5 μm, and stained with haematoxylin and eosin. Slides were scanned using a Zeiss Axioscan Z1 slide scanner. HALO Image Analysis Software (Indica Labs) was used to quantify vacuolar area and number of nuclei within BAT sections.

### Serum biochemistry

Serum triglycerides and free fatty acids were measured using a Dimension RXL analyser (Siemens Healthcare) or the Roche Free Fatty Acid Kit (half-micro test, 11383175001), respectively.

### Glucose tolerance test

For the intraperitoneal glucose tolerance test, mice were fasted overnight for 16 h with free access to water. Glucose (2 g/kg body weight) was administered intraperitoneally, and blood glucose was measured from the tail tip at indicated time points using a glucometer.

### Norepinephrine-stimulated energy expenditure

Energy expenditure was assessed by indirect calorimetry in a temperature-controlled Oxymax chamber (2.7 L) maintained at 30°C, with airflow set to 400 mL/min. Oxygen consumption (VO) and carbon dioxide production (VCO) were measured every 4 min using a custom-built monitoring system (Creative Scientific), and EE was calculated using a modified Weir equation (EE J/min= 15.818xVO2 (ml/min) + 5.176*VCO2 (ml/min). Mice were housed at 10°C for 3 days prior to the experiment. Following pentobarbital anaesthesia (60 mg/kg), mice were allowed to reach a steady-state baseline (three consecutive stable measurements, average considered baseline). Mice were then administered norepinephrine bitartrate (1 mg/kg) together with pentobarbital (30 mg/kg) subcutaneously and returned to the calorimetry chamber. EE was recorded for a further 28 min. Maximum thermogenic capacity was defined as the mean of the three highest EE measurements following norepinephrine administration.

### Gene silencing

Gene silencing was performed using Silencer Select siRNAs (s1500, s1501 and s1502, 10 nM) and Lipofectamine RNAiMAX (Cat# 13778150, Thermo Fisher Scientific) according to the manufacturer’s instructions. Briefly, cells were transfected in antibiotic-free medium with siRNA–RNAiMAX complexes prepared in Opti-MEM (Cat# 31985070, Thermo Fisher Scientific). Following 10–15 min incubation at room temperature, complexes were added dropwise to cells and medium was replaced after 16 h.

### RNA extraction and bulk RNA-seq

RNA was extracted using Qiagen RNeasy kit (74106), following manufacturer’s instructions. Total RNA (>200 ng) was submitted to Novogene for bulk RNA sequencing using standard library preparation and sequencing workflows. Samples were sequenced on an Illumina platform with paired-end 150 bp reads (PE150), generating approximately 6 Gb of raw data per sample.

Raw sequencing reads in FASTQ format were assessed using FastQC and transcript abundance quantified using Salmon. RNA-seq abundance estimates were imported into R using “tximport” (v1.38.2). Differential gene expression analysis was performed using “DESeq2” (v1.50.2), with p-values adjusted using the Benjamini–Hochberg procedure to control the false discovery rate (FDR). Pathway enrichment analysis was conducted using “fgsea” (v1.36.2) based on ranked gene lists derived from differential expression results. Transcription factor activity was inferred using "viper” (v1.44.0) and the “CollecTri” repository from “OmnipathR” (v3.18.3) to estimate regulator activity from gene expression signatures. All analyses were performed in R (v4.5.0).

### Statistical analysis

The number of animals or independent experiments is indicated in the corresponding figure legends. Data are presented as mean ± SEM, unless otherwise specified; for violin plots, centre lines indicate the median. Statistical analyses were performed using GraphPad Prism 11, and specific tests indicated in the figure legends. Comparisons among more than two groups were performed using one-way ANOVA followed by Dunnett’s multiple-comparisons test. Analyses including more than one factor were performed using two-way ANOVA followed by Dunnett’s multiple-comparisons test. Repeated-measures versions of these tests were used where appropriate. Statistical significance was defined as P < 0.05.

## Data availability

Human PVAT snRNA-seq data from three male patients undergoing coronary artery bypass surgery, as described in Angueira et al. ^24^, was downloaded from the Gene Expression Omnibus (GEO) database (GSE164528; samples GSM5068996, GSM5068997, GSM5068998). All written code is available at https://doi.org/10.5281/zenodo.22811626.

The bulk RNA-seq data reported in this paper have been deposited in GEO under accession number GSE342095.

## Acknowledgments

We thank **Nicholas M. Katritsis** for helpful discussions and for critically reviewing and revising the manuscript. We thank the Histopathology, Imaging, and Disease Model Cores and the Core Biochemistry Assay Laboratory at the Wellcome-MRC Institute of Metabolic Science Metabolic Research Laboratories for their help with this work.

## Author contributions

S.R.-F. and A.V.-P. conceived the original hypothesis. F.B. and N.H. designed and supervised the bioinformatics analyses. H.M.L.B. and F.B. performed the bioinformatics analyses. S.R.-F. and M.M. performed the bulk RNA-seq bioinformatics analysis. S.R.-F., N.G., M.C. and M.M. performed *in vivo* experiments. S.R.-F. designed, performed and supervised the *in vitro* and *in vivo* experiments. S.R.-F., I.D.M. and T.L.S. performed *in vitro* experiments. P.P. generated and provided data. H.M.L.B., F.B. and S.R.-F. wrote the manuscript. A.V.-P. and N.H. edited and discussed the manuscript. All authors approved this publication.

