## Supplementary figures for "An amyloid-β/APP regulatory layer in human brown adipocyte recruitment identifies repurposable drugs for thermogenic fat expansion"

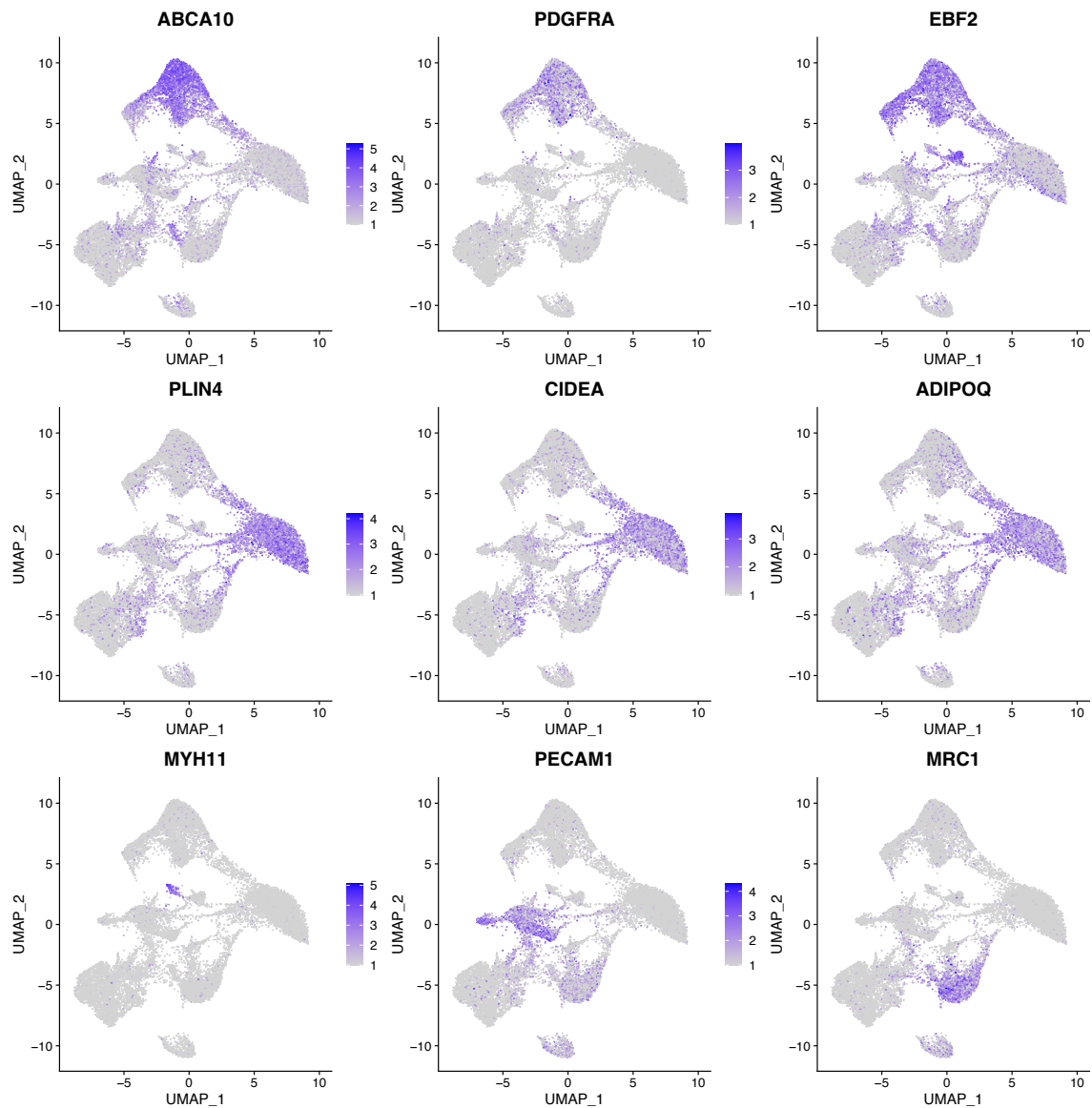

**Figure S1 | Expression patterns of marker genes across UMAP clusters were used for cell type identification.** Feature plots displaying expression of key marker genes overlaid on UMAP coordinates from Fig. 2A. Brown adipocyte lineage markers include *ABCA10*, *PDGFRA*, and *EBF2* (preadipocyte stages), *PLIN4*, *CIDEA*, and *ADIPOQ* (mature brown adipocytes). Non-adipocyte markers include *MYH11* (vascular smooth muscle cells), *PECAM1* (endothelial cells), and *MRC1* (macrophages). These expression patterns confirm the cluster annotations in Fig. 2B and guide downstream pathway enrichment analyses to characterise brown adipocyte differentiation. Colour intensity indicates expression level (purple = high, grey = low/absent). n = 18,780 nuclei from three male donors. UMAP, Uniform Manifold Approximation and Projection.

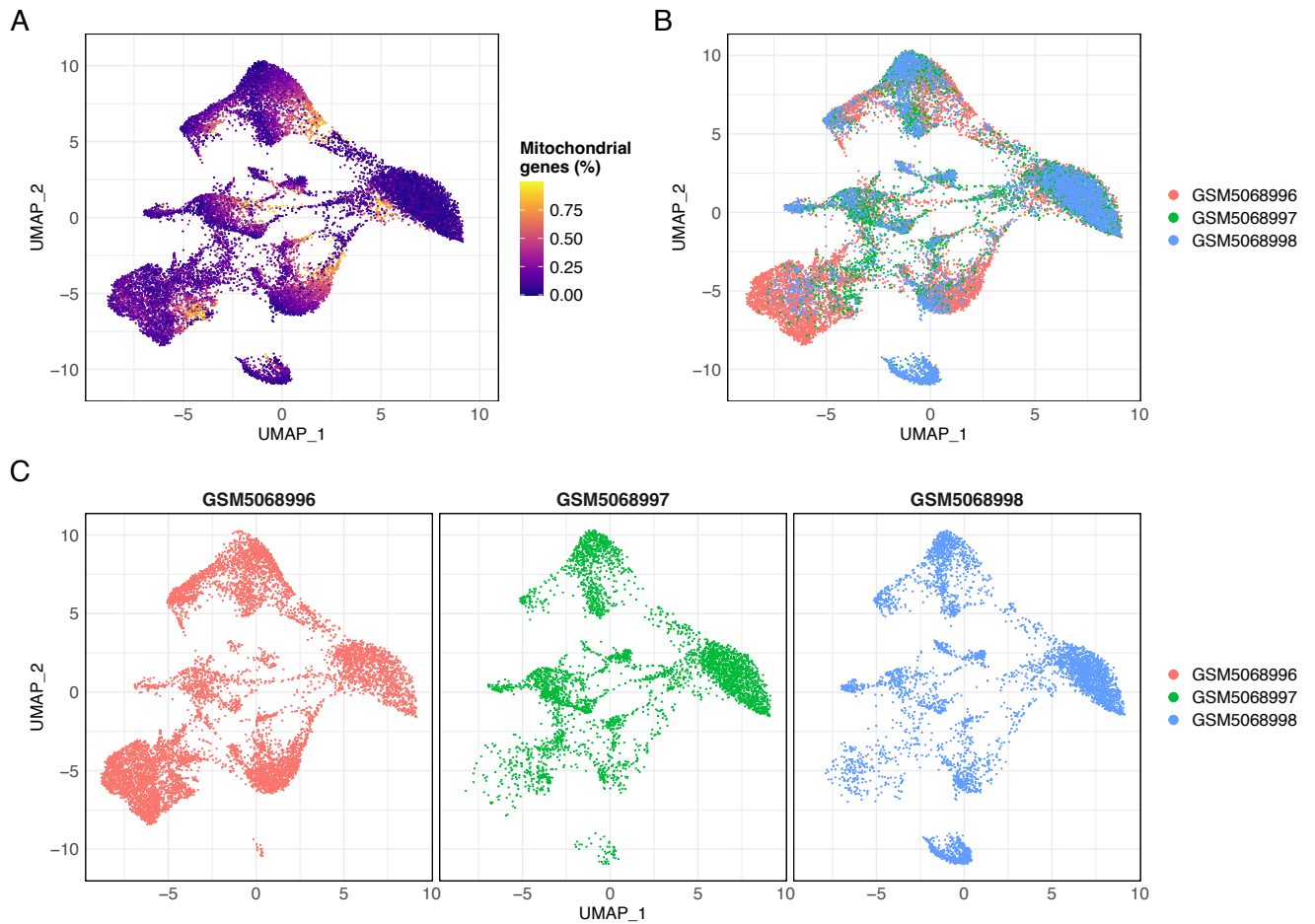

**Figure S2 | UMAP quality control and sample distribution assessment. (A)** Mitochondrial gene content per nucleus; all retained nuclei fall below the 1% threshold applied during quality control filtering. **(B)** Sample type distribution across UMAP coordinates showing successful integration of three samples (n = 18,780 nuclei from three male donors). **(C)** UMAP visualisation split by individual donor, showing comparable representation of each cluster across the three donors and no donor-restricted cluster. UMAP, Uniform Manifold Approximation and Projection.

A

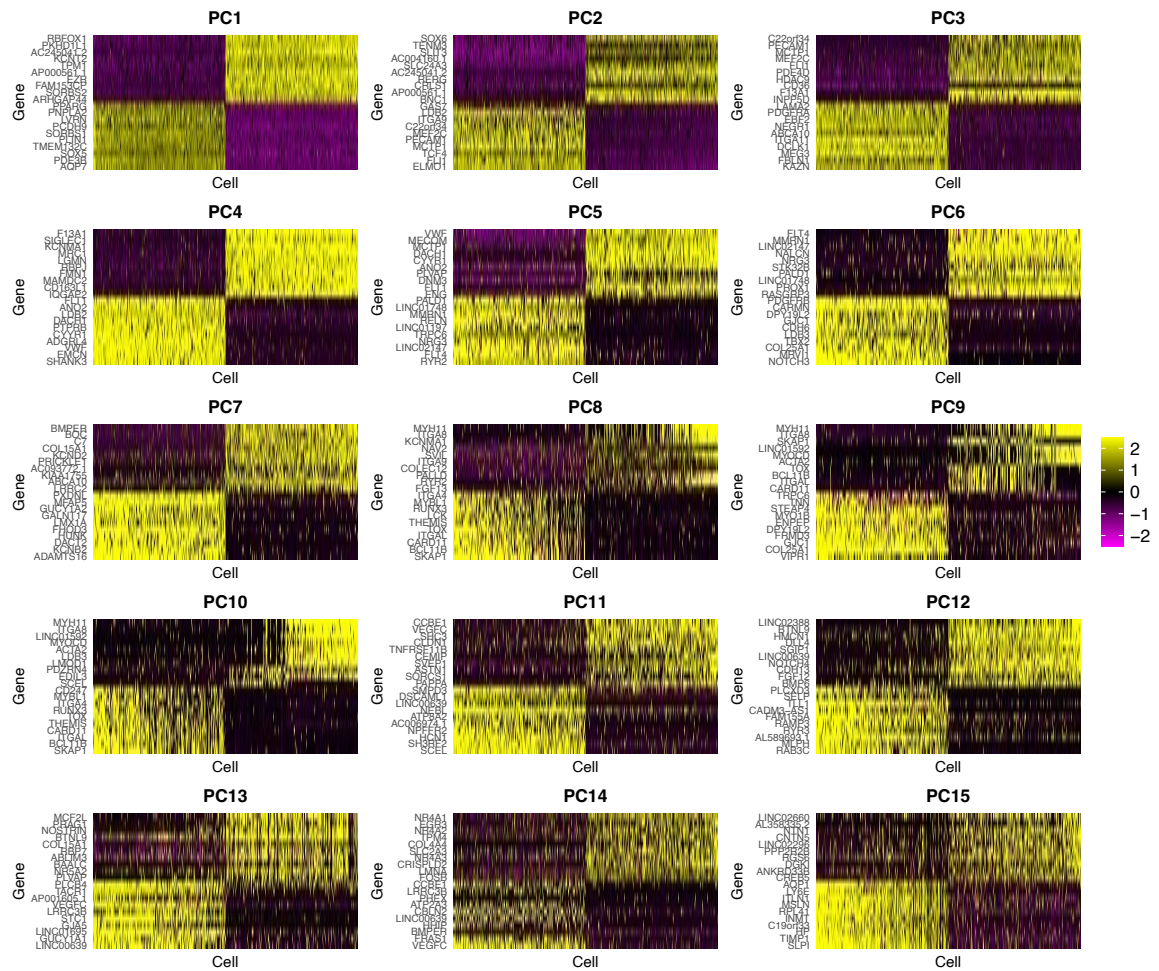

B

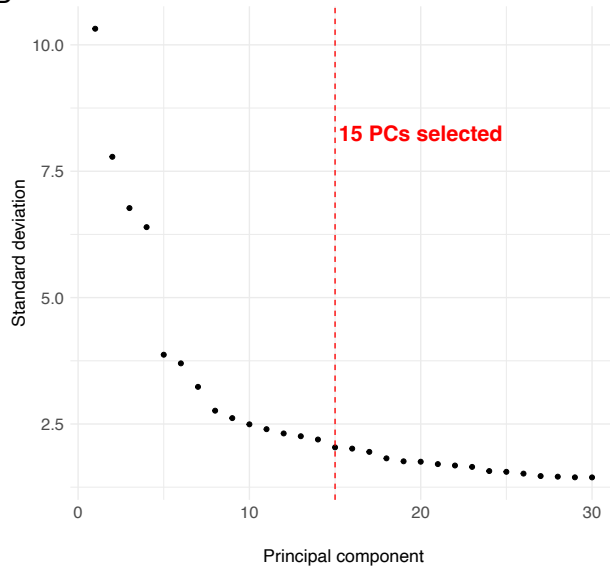

C

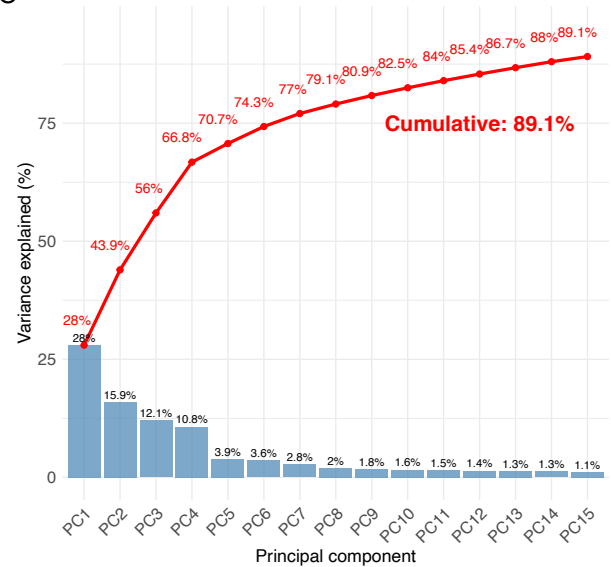

**Figure S3 | Principal component analysis and variance decomposition of integrated snRNA-seq data. (A)** Heatmaps showing the top 20 genes with the highest loadings for principal components (PCs) 1-15, displayed across 500 randomly sampled nuclei ($n = 18,780$ nuclei in total). Yellow indicates high expression; purple indicates low expression. Each PC captures distinct gene expression patterns that drive cellular heterogeneity. **(B)** Elbow plot showing the standard deviation explained by each principal component (y-axis) versus PC number (x-axis).

The inflection point around PC15 (red dashed line) guided the selection of 15 PCs for downstream dimensionality reduction. **(C)** Variance decomposition by principal component, showing individual variance contribution by each PC (blue bars) and cumulative variance (red line). The first 15 PCs collectively capture 89.1% of total variance in the dataset, providing sufficient dimensionality reduction for downstream UMAP analysis. UMAP, Uniform Manifold Approximation and Projection.

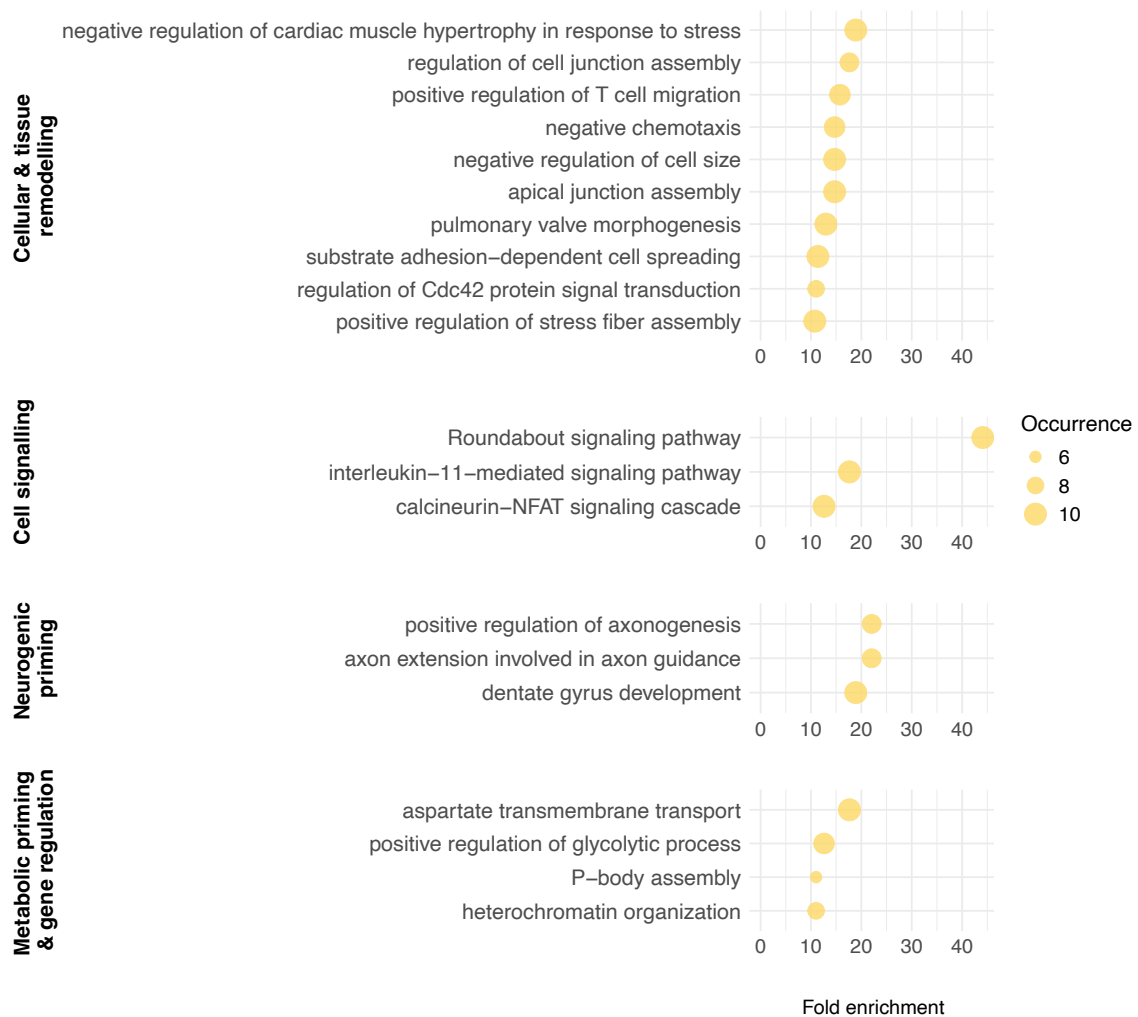

Figure S4 | **Pathway enrichment analysis in early- to intermediate-stage preadipocyte transition reveals neurogenic signalling mechanisms alongside canonical remodelling processes.** We grouped enriched pathways into biological themes: cellular and tissue remodelling, cell signalling, neurogenic priming, and metabolic priming and gene regulation. Fold enrichment refers to the extent of a pathway's statistical overrepresentation in a given differentiation step without inferring directionality (e.g. up- or downregulation in one adipocyte stage or another). Occurrence reflects enrichment frequency over 10 iterations. Italicised text indicates pathway names.

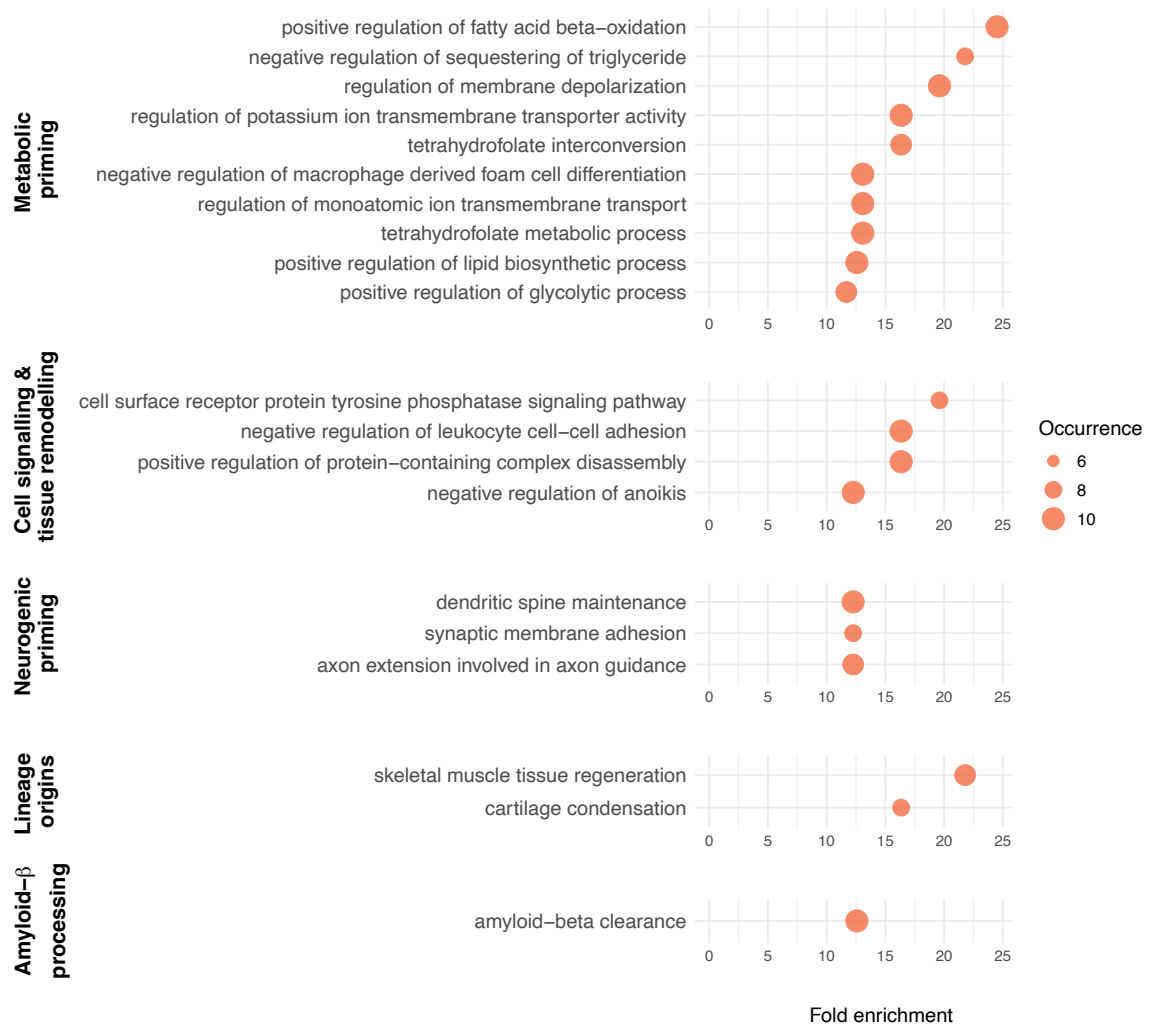

Figure S5 | **Pathway enrichment analysis in intermediate preadipocytes to transitional adipocytes transition reveals increased metabolic priming, sustained neurogenic priming, and emerging amyloid-β clearance mechanisms.** We grouped enriched pathways into biological themes: metabolic priming, cell signalling and tissue remodelling, neurogenic priming, lineage origins, and amyloid-β processing. Fold enrichment refers to the extent of a pathway's statistical overrepresentation in a given differentiation step without inferring directionality (e.g. up- or downregulation in one adipocyte stage or another). Occurrence reflects enrichment frequency over 10 iterations. Italicised text indicates pathway names.

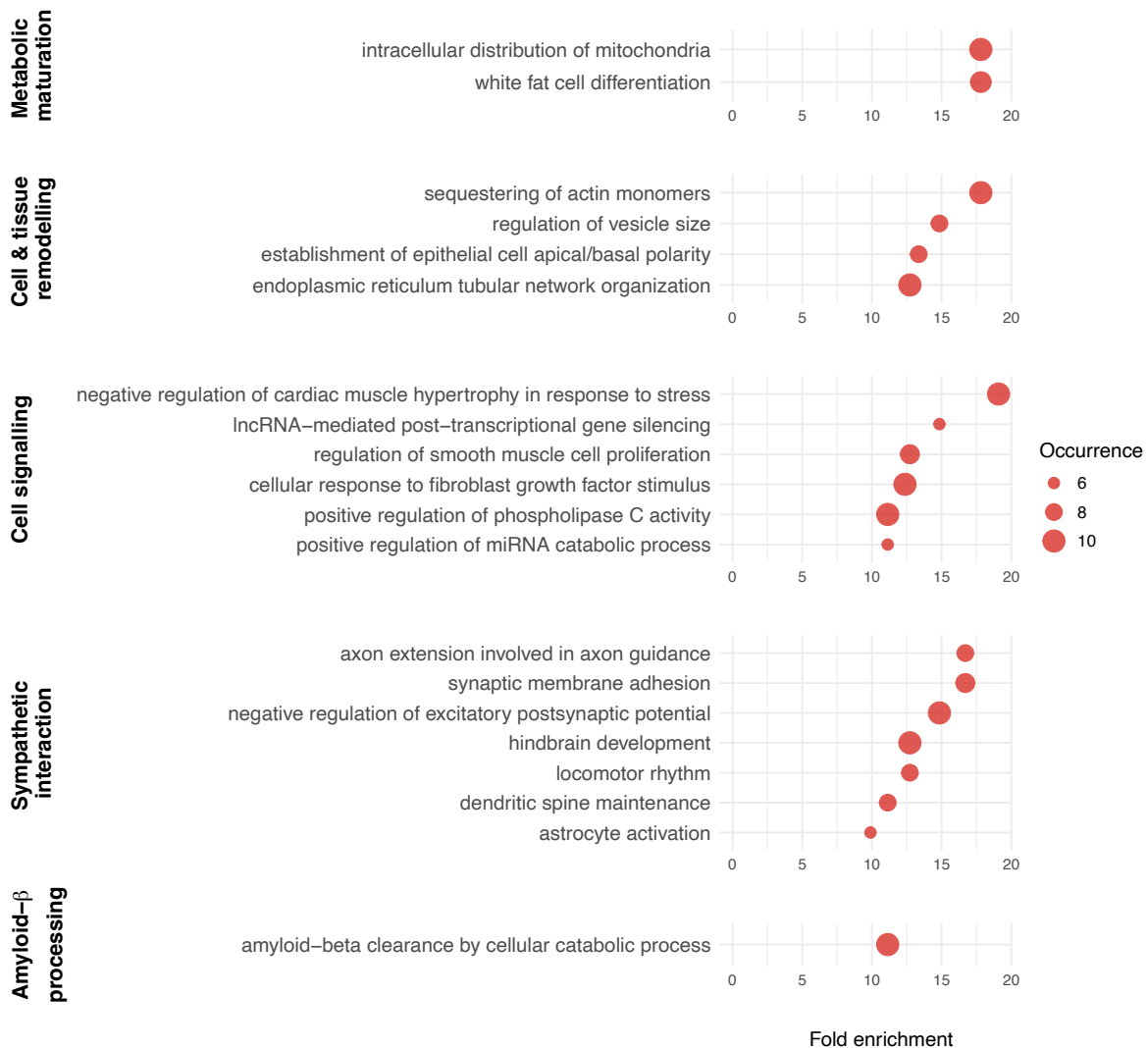

Figure S6 | **Pathway enrichment analysis in transitional to mature adipocytes transition shows metabolic maturation, sustained sympathetic interaction, and maintained amyloid-β processing capacity.** Enriched pathways were grouped into biological themes: metabolic maturation, cell and tissue remodelling, cell signalling, sympathetic interaction, and amyloid-β processing. Fold enrichment refers to the extent of a pathway's statistical overrepresentation in a given differentiation step without inferring directionality (e.g. up- or downregulation in one adipocyte stage or another). Occurrence reflects enrichment frequency over 10 iterations. Italicised text indicates pathway names.

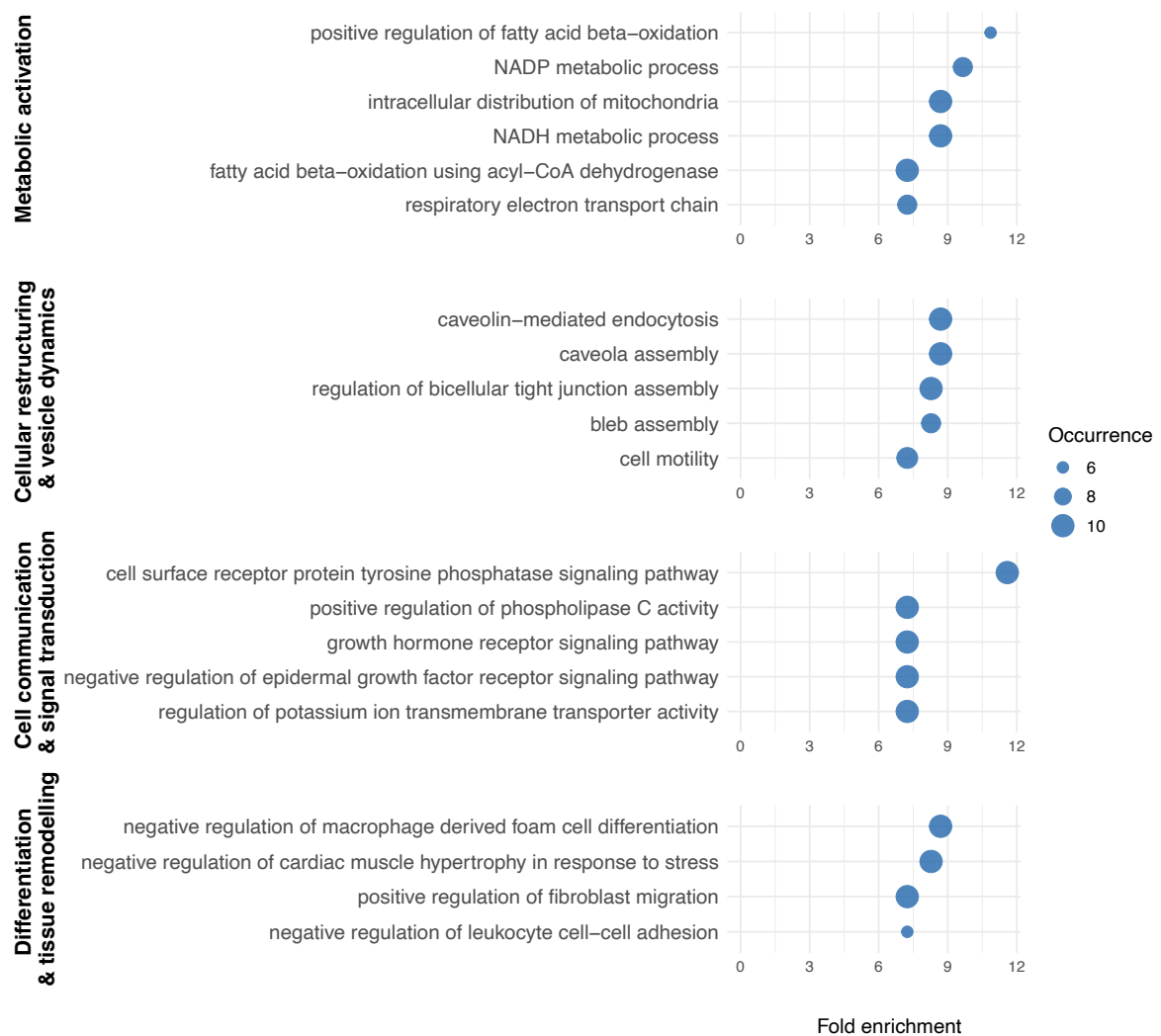

Figure S7 | **Pathway enrichment analysis of full brown adipocyte differentiation (early preadipocytes to mature adipocytes) shows metabolic activation, cellular restructuring and cell signalling.** Enriched pathways were grouped into biological themes: metabolic activation, cellular restructuring and vesicle dynamics, cell communication and signal transduction, and differentiation and tissue remodelling. Fold enrichment refers to the extent of a pathway's statistical overrepresentation in a given differentiation step without inferring directionality (e.g. up- or downregulation in one adipocyte stage or another). Occurrence reflects enrichment frequency over 10 iterations. *Italicised text indicates pathway names.*

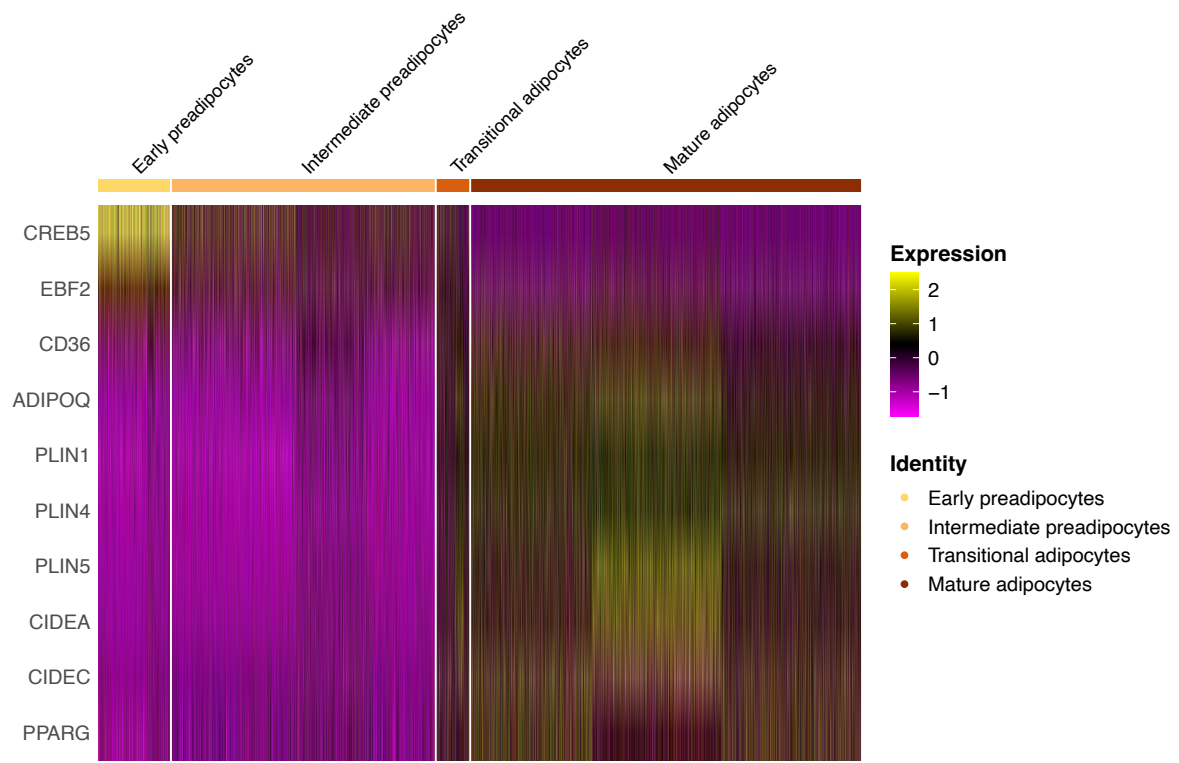

Figure S8 | **Marker gene expression shows stage-specific dynamics during brown adipocyte differentiation.** Heatmap shows z-scored expression of ten marker genes across cells grouped by stage (early preadipocytes, intermediate preadipocytes, transitional adipocytes, and mature adipocytes). Early-stage markers (*CREB5*, *EBF2*) peak early, while late-stage markers (*CD36*, *ADIPOQ*, *PLIN1*, *PLIN4*, *PLIN5*, *CIDEA*, *CIDEA*, *PPARG*) increase progressively.

### A. PPI network construction

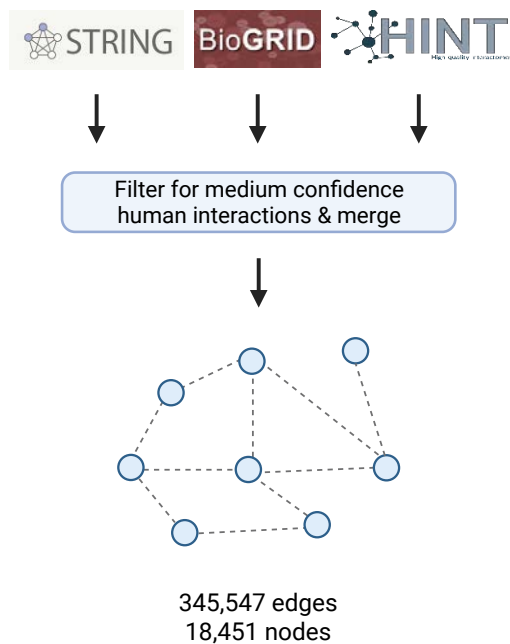

### B. DPI network construction

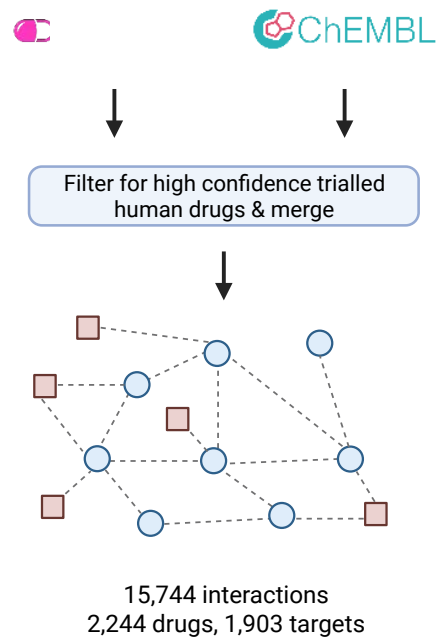

### C. Drug proximity analysis

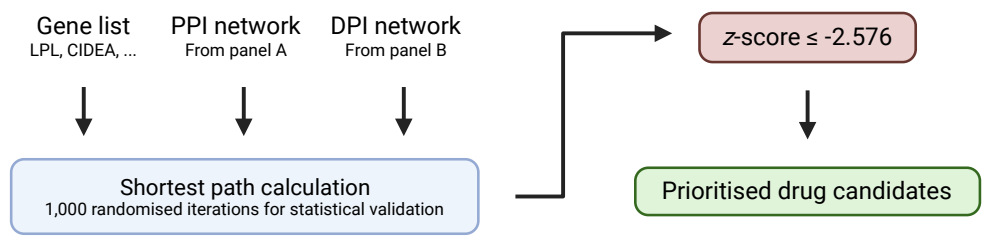

**Figure S9 | PPI network-based drug prioritisation methodology for identifying therapeutic compounds targeting brown adipocyte differentiation pathways.** (A) Protein-protein interaction (PPI) network construction by merging STRING, BioGRID and HINT, retaining medium-confidence ($\geq 0.4$) STRING interactions and multi-validated physical BioGRID interactions, yielding 345,547 edges and 18,451 nodes. (C) Drug-protein interaction (DPI) network construction from DrugBank and ChEMBL databases, filtering for high-confidence human drugs and targets, resulting in 15,744 unique interactions between 2,244 drugs and 1,903 protein targets. (B) Proximity algorithm workflow combining the input gene list with the PPI and DPI networks to compute the average shortest path length from drug targets to input genes. Significance was assessed against 1,000 degree-matched randomisations, and drugs with $z \leq -2.576$ were retained.

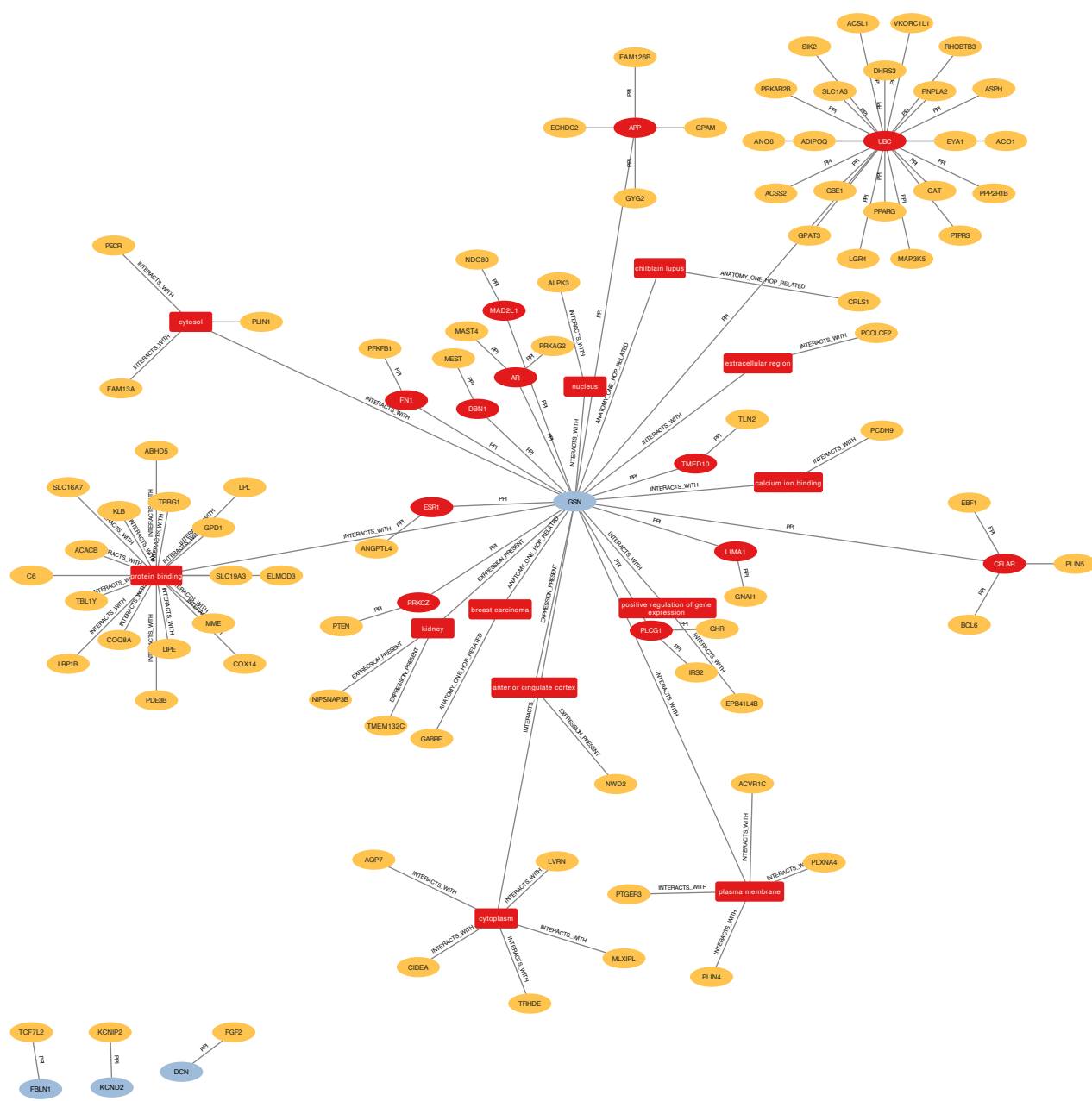

**Figure S10 | Knowledge graph traversal between markers of preadipocytes and those of adipocytes reveals novel gene modulators in the hidden layer.** Traversing the graph starting from the differentially expressed genes in preadipocytes and terminating at the differentially expressed genes in adipocytes reveals a hidden layer of regulatory genes not apparent from the transcriptomic analysis alone; those discussed in the main text are MAD2L1, PRKCZ, CFLAR, LIMA1, DBN1, TMED10, ATF3, FN1, AR, ESR1 and APP. Note that edges between resultant nodes in the graph are hidden for clarity.

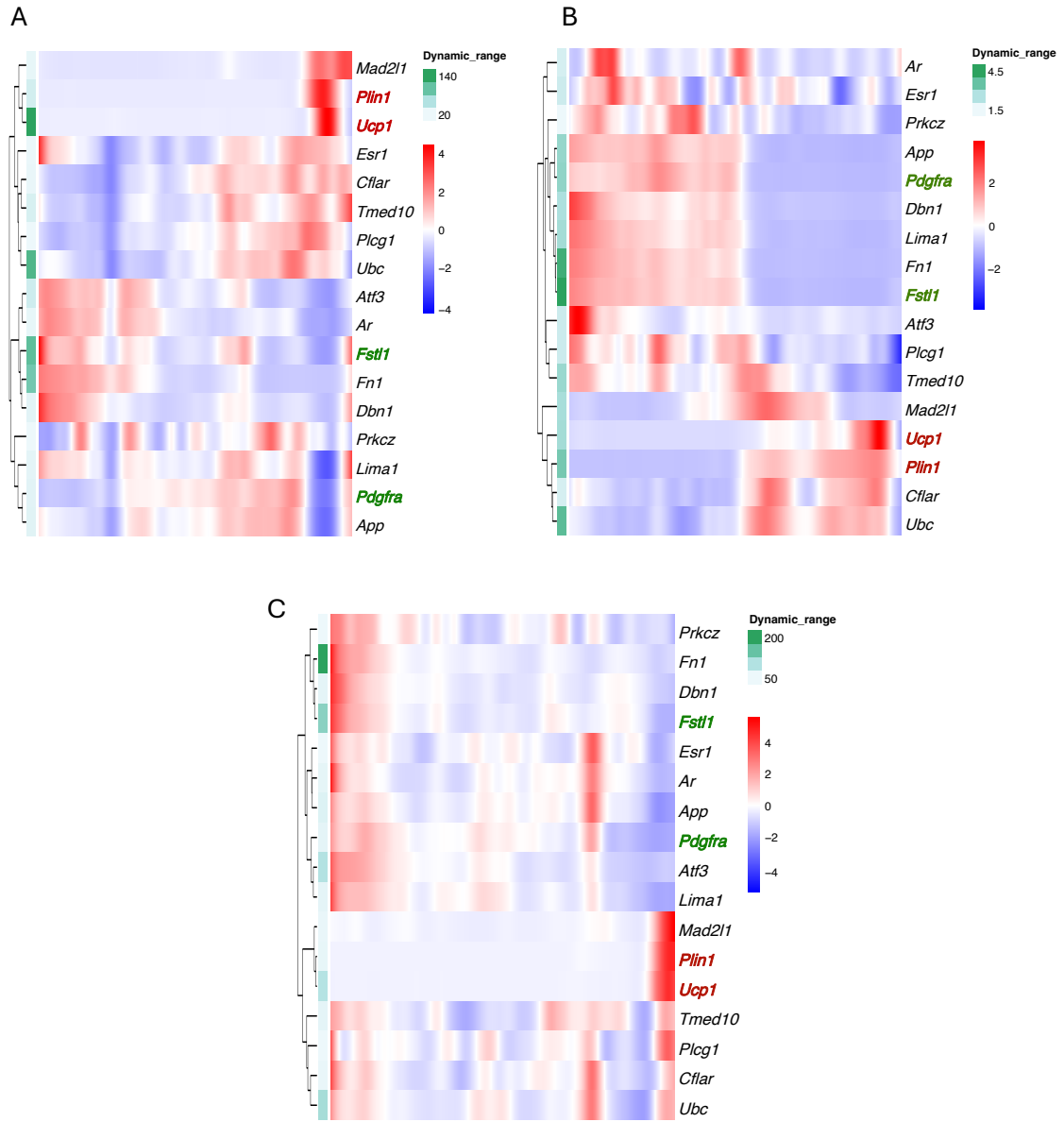

Figure S11 | **Heatmaps showing gene expression across brown adipocyte differentiation in the hidden layer from preadipocyte to adipocyte markers. (A-C).** Hierarchical clustering was applied to identify clustered expression patterns of key markers of preadipocytes (green), markers of adipocytes (red), and genes identified in the hidden layer from KG-based network analyses. Nuclei are ordered in pseudotime from trajectory analysis of published mouse datasets: (A) BAT recruitment (GSE207706), (B) BAT embryonic development (GSE185623), and (C) perinatal differentiation in para-aortic perivascular adipose tissue (GSE164528).

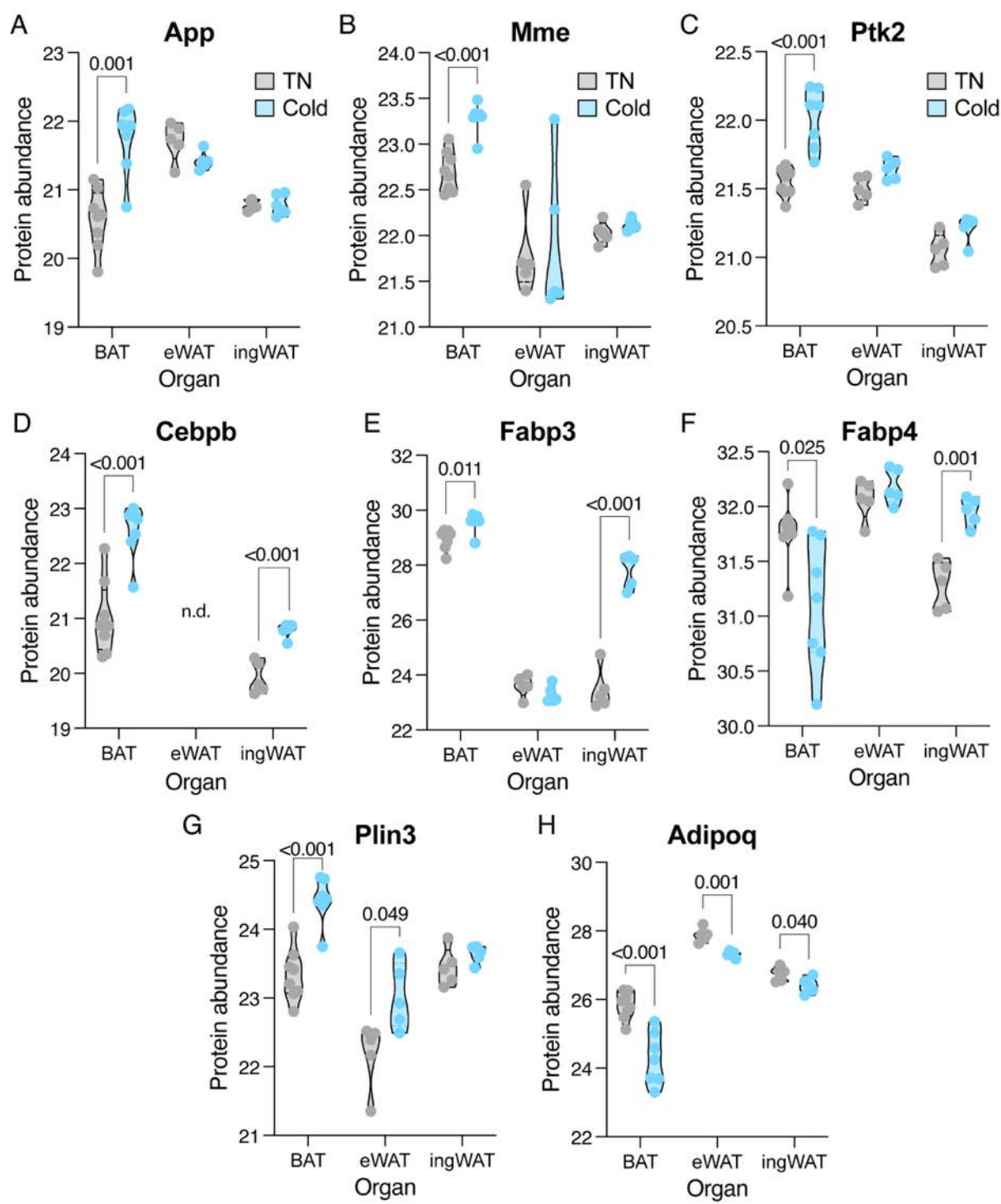

Figure S12 | **Network-designated nodes increase in abundance in brown, but not white, adipose depots during cold-induced recruitment.** (A–H) Abundance of the indicated proteins in interscapular BAT, ingWAT and eWAT of mice exposed to cold, measured by mass spectrometry. Reanalysis of published data (Cutler et al., Science Advances, 2025); cold exposure duration as reported in that study. eWAT, epididymal white adipose tissue; ingWAT, inguinal white adipose tissue.

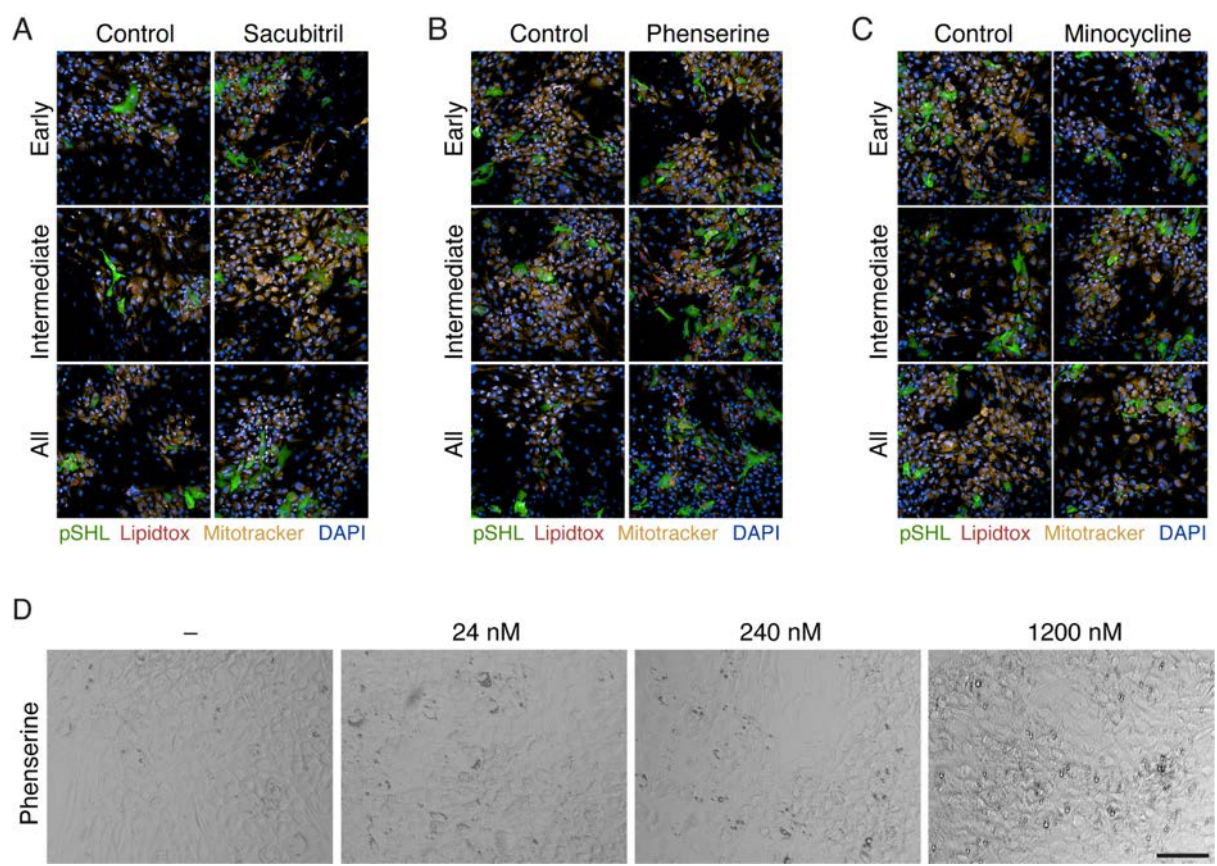

Figure S13 | ***In vitro* validation identified drugs enhancing brown adipocyte differentiation.** **(A-C)** Representative confocal images of cells differentiated in the presence of sacubitril (250 nM) (A), phenserine (1200 nM) (B) or minocycline (500 nM) (C) and examined on day 18 of differentiation under basal conditions. Representative fields were selected automatically based on quantitative analysis to identify images closest to the mean across all channels—pHSL, phosphorylated HSL (Ser660). **(D)** Representative images of brown adipocytes on day 10 of differentiation treated with the indicated concentration of phenserine throughout the differentiation process. Scale bar: 100 $\mu\text{m}$.

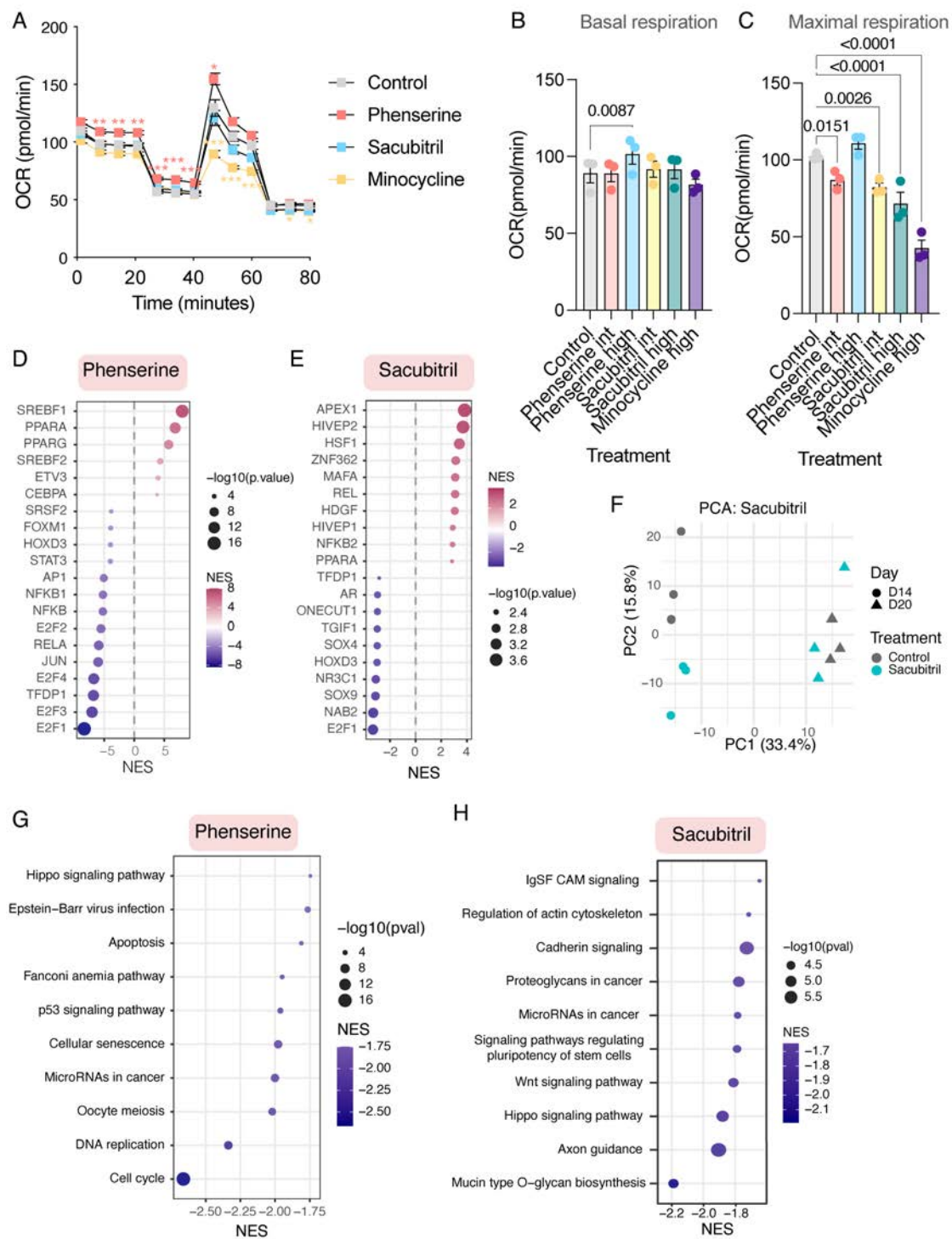

**Figure S14 | Distinct phenotype and molecular profile of positive hits. (A-C)** Representative Seahorse Mito Stress Test profile (A) and quantification of basal (B) and maximal respiration (C) in differentiated hTERT-BA cells treated with the indicated compounds and concentrations throughout the differentiation process. Repeated-measures two-way ANOVA was used to assess treatment, dose, and treatment × dose interaction effects, followed by Dunnett's post hoc test comparing each condition with the corresponding untreated control. Data are presented as mean ± SEM from three independent experiments and are normalised to the untreated control. **(D, E)** Upstream regulator analysis of bulk RNA-seq from fully differentiated cells treated

throughout differentiation with phenserine (D) or sacubitril (E). **(F)** Principal component analysis of cells treated with sacubitril throughout differentiation, with samples collected on days 14 and 20 of differentiation. **(G, H)** Top downregulated pathways identified by pathway enrichment analysis of the same transcriptomic dataset shown in D, E. OCR, oxygen consumption rate; NES, normalised enrichment score; PC, principal component.

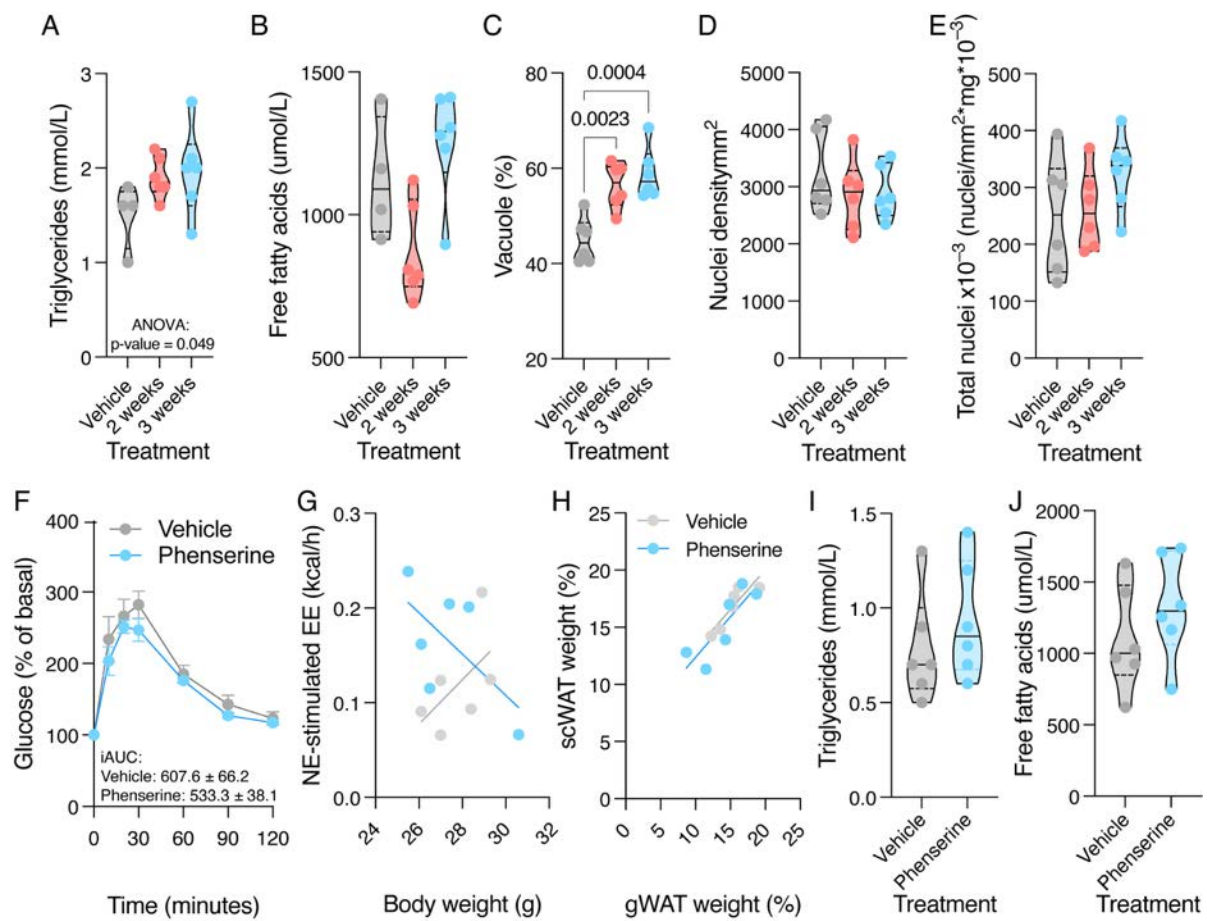

**Figure S15 | Phenserine alters BAT lipid handling and the BAT-WAT relationship *in vivo*. (A-B)** Terminal serum triglycerides (A) and free fatty acids (B) in vehicle- and phenserine-treated mice at rest. **(C-E)** Quantification of BAT lipid vacuole area (C, % tissue area), nuclei density (D, nuclei/mm<sup>2</sup>), and estimated total nuclei number (E). **(F)** Intraperitoneal glucose tolerance test in mice treated with vehicle or phenserine for 3 weeks. **(G-H)** Linear regression between the indicated parameters in mice treated for 3 weeks, followed by 3 days of cold exposure at 10°C and norepinephrine stimulation. **(I-J)** Terminal serum triglycerides (I) and free fatty acids (J) in mice treated as described for G- H. Data are shown as violin plots, with the central line indicating the median, or as mean ± SEM. N = 6 male C57BL/6J mice per group in each experiment. One-way ANOVA followed by Dunnett's post hoc test was used where three or more groups were compared; two-group comparisons were made by Student's *t*-test (A-E, F (iAUC), I-J). Slopes and elevations of the regressions in G and H were compared by analysis of covariance.

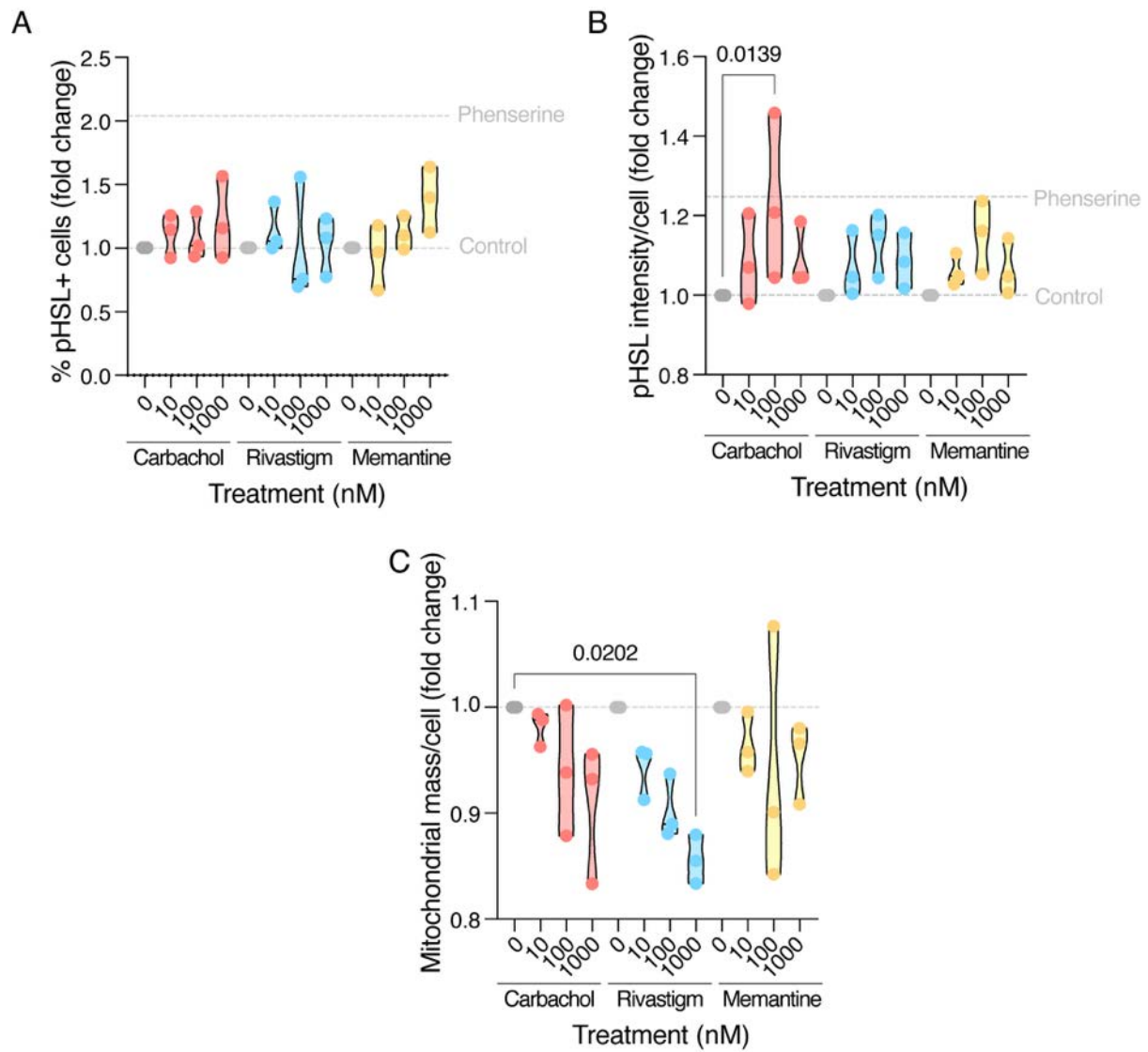

Figure S16 | **Cholinergic drugs could not recapitulate phenserine treatment effects. (A–C).** Quantification of the proportion of cells positive for HSL phosphorylation (A), average phosphoHSL signal intensity per cell (B) and average MitoTracker signal per cell (C) in hTERT-BA cells differentiated in the presence of carbachol, rivastigmine or memantine at the indicated concentrations, as assessed by high-content imaging analysis. Repeated measures two-way ANOVA was used to test for treatment, dose, and interaction effects, followed by Dunnett's post hoc test comparing each treatment with the corresponding control. Data from three independent experiments are shown as violin plots, normalised to the untreated control. Values obtained from phenserine-treated cells in the same experiments are indicated by dashed lines.
